# *Dandelion* utilizes single cell adaptive immune receptor repertoire to explore lymphocyte developmental origins

**DOI:** 10.1101/2022.11.18.517068

**Authors:** Chenqu Suo, Krzysztof Polanski, Emma Dann, Rik G.H. Lindeboom, Roser Vilarrasa-Blasi, Roser Vento-Tormo, Muzlifah Haniffa, Kerstin B. Meyer, Lisa M. Dratva, Zewen Kelvin Tuong, Menna R. Clatworthy, Sarah A. Teichmann

**Author notes:** Corresponding authors. (Z.K.T.), (M.R.C.), (S.A.T.). These authors contributed equally to this work (Chenqu Suo, Krzysztof Polanski). These senior authors contributed equally to this work (Zewen Kelvin Tuong, Menna R. Clatworthy, Sarah A. Teichmann).

## Abstract

Assessment of single-cell gene expression (scRNA-seq) and adaptive immune receptor sequencing (scVDJ-seq) has been invaluable in studying lymphocyte biology. Here, we introduce *Dandelion*, a computational pipeline for scVDJ-seq analysis. It enables the application of standard V(D)J analysis workflows to single-cell datasets, delivering improved V(D)J contig annotation and the identification of non-productive and partially spliced contigs. We devised a novel strategy to create an adaptive immune receptor feature space that can be used for both differential V(D)J usage analysis and pseudotime trajectory inference. The application of *Dandelion* improved the alignment of human thymic development trajectories of double positive T cells to mature single-positive CD4/CD8 T cells, with important new predictions of factors regulating lineage commitment. *Dandelion* analysis of other cell compartments provided novel insights into the origins of human B1 cells and ILC/NK cell development, illustrating the power of our approach. *Dandelion* is an open access resource (https://www.github.com/zktuong/dandelion) that will enable future discoveries.

## Introduction

Recent developments in single-cell genomics have significantly advanced our understanding of human immunology^1, 2^. Paired adaptive immune receptor (AIR) sequencing with mRNA expression in the same cell allows for direct linkage of AIR repertoire with cellular phenotypes, and has proven to be a powerful tool in understanding lymphocyte development and function in healthy and disease contexts^3–6^.

Multi-omics analysis leverages data from different modalities and has been successfully applied in recent years to study cellular biology at an unprecedented resolution. Examples include integration of paired single-cell RNA sequencing (scRNA-seq) and Assay for Transposase-Accessible Chromatin with high-throughput sequencing (ATAC-seq) data or Cellular Indexing of Transcriptomes and Epitopes by Sequencing (CITE-seq) data^7, 8^.

However, unlike many other sequencing modalities, which largely consist of continuous data, AIR repertoire sequencing data are a mixture of categorical and continuous data which pose additional challenges for integration. AIR data consist of annotations of variable (V), diversity (D) and joining (J) genes, which are recombined and selected during B/T cell development^9^. The Adaptive Immune Receptor Repertoire (AIRR) community was formed in 2015 to help address the issues and challenges related to the curation and analysis of AIR data generated with high throughput sequencing technologies^10–12^. This has led to the standardization of repertoire data representation across various modes of AIR data, including single-cell V(D)J sequencing data. There are established options and packages that can deal with single-cell AIR repertoire data and they provide a variety of methods for downstream analyses (non-exhaustive list of some popular tools shown in **Supplementary Fig. 1**). The functions include re-annotation of genes in AIR contigs, quality control checks, matching contigs to cells, clonotype definition, mutation quantification and diversity estimation and many more (**Supplementary Fig. 1**). The single-cell AIR softwares are often designed to interact with a single-cell gene expression software package of choice, e.g. *scirpy*^13^ with *scanpy*^14^ and *scRepertoire*^15^ with *Seurat*^16^, providing valuable visualization options. There are also tools for predicting antigen specificity of T cell receptors (TCRs) (e.g. *TcellMatch*^17^), annotating TCRs that recognize known epitopes (e.g. Platypus^18^, Immunarch^19^) and extraction of significant motifs and motif groups (e.g. *ALICE*^20^). There have also been developments in joint-embedding of single-cell gene expression and AIR complementarity-determining region 3 (CDR3) sequences, which have been used for correlating TCRs with T cell transcriptomes (e.g. *CoNGA*^21^, *mvTCR*^22^). There remains opportunities for new methods to realize the full potential of paired scRNA-seq and scVDJ-seq data.

To that end, we developed *Dandelion*, a holistic analysis framework within the context of single-cell lymphocyte biology. It offers a B cell receptor (BCR) and TCR contig annotation pipeline, integrative analysis with single cell RNA-seq data and a novel V(D)J feature space for differential V(D)J usage and pseudotime trajectory inference. Here, using two immune development datasets, we showcase how *Dandelion* can be applied to improve alignment of cells along the double positive (DP) T cell to mature T cell development trajectory, and provide novel insights into human B1 cell origin and innate lymphoid cell (ILC) and natural killer (NK) cell development.

## Results

### *Dandelion* enables holistic scVDJ-seq analysis

As *Dandelion* operates on the AIRR data format, it has high interoperability with existing tools in the AIRR community^13, 23^ and can serve as a bridge between these tools and single-cell gene expression analysis software ecosystem e.g. *scverse*^14, 24^ (**Fig. 1a**). *Dandelion* has also been certified by the AIRR Software Working Group to be compliant with the software standards that encourage collaboration and reproducibility.

**Fig. 1.**
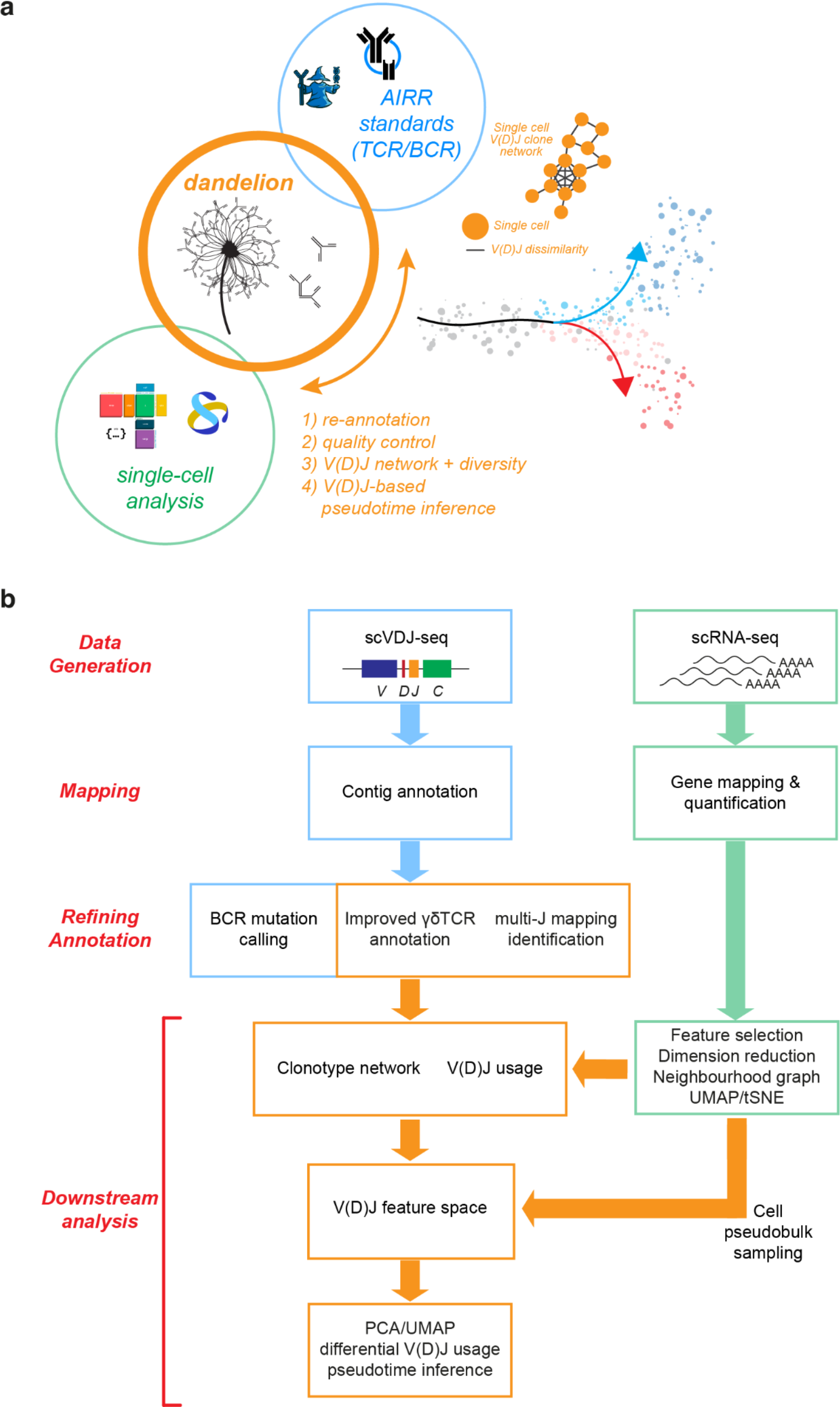
Holistic scVDJ-seq analysis pipeline. a, Schematic illustration showing that *Dandelion* bridges methods from single-cell V(D)J workflows such as AIRR standards and the single-cell gene expression analysis software, and combines with them additional novel methods of its own to create a holistic pipeline for analysis. **b,** Schematic illustration of the *Dandelion* workflow. Paired single-cell gene expression (scRNA-seq) and AIR repertoire (scVDJ-seq) data is generated, followed by mapping of the sequencing reads. From the mapped results, *Dandelion* provides refined contig annotations with BCR mutation calling, improved γδTCR mapping and identification of multi-J mapping contigs. It also provides downstream analysis after integration with scRNA-seq results. Apart from allowing the users to explore clonotype networks and V(D)J usage, *Dandelion* also supports building a V(D)J feature space on pseudo-bulked cells, that can be used for differential V(D)J usage and pseudotime inference. Additional unique features provided by *Dandelion* are boxed in orange.

*Dandelion* can be used to analyze single-cell BCR, αβTCR and γδTCR data, allowing for BCR mutation calling, improved γδTCR mapping, extraction of both productive and non-productive V(D)J contigs and identification of unspliced J gene alignments (‘multi-J mapping’) (**Fig. 1b**). *Dandelion* then performs quality control checks, clonotype calling and clonotype network generation for downstream analyses, and is designed to work with any AIRR formatted input or 10X Genomics’ *cellranger vdj* output. A main novel feature of *Dandelion* is the creation of a ‘V(D)J feature space’ that can be used to visualize TCR/BCR usage across cell pseudo-bulks or neighborhoods, perform differential V(D)J usage analysis and pseudotime trajectory inference. A summary list of features of *Dandelion* and a non-exhaustive list of other existing pipelines is shown in **Supplementary Fig. 1**. A subset of the functionalities of *Dandelion* was previously applied to a large COVID-19 study^4^ which showcased its network-based repertoire diversity analysis method.

### *Dandelion* provides a streamlined contig re-annotations pipeline

For optional re-annotation of contigs, *Dandelion* expects 10X Genomics’ *cellranger vdj* output files, specifically the contig annotation spreadsheet and fasta file (e.g. *all_contig_annotations.csv* and *all_contig.fasta*).

Similar to *Change-O*^23^, *Dandelion* re-annotates V(D)J contigs using *igblastn*^25^ with reference sequences contained in the international ImMunoGeneTics information system (IMGT) database^26^. The individual contigs are then checked with *blastn* for the D and J gene separately, using the same settings as per *igblastn*^25^. The additional *blastn* step allows us to: i) apply an e-value cut off for D and J calls to ensure only high confidence calls are retained; ii) identify multi-J mapping contigs (see below); and iii) recover contigs without V gene calls (removed by *igblastn*). We packaged this pre-processing workflow into a single-line command implemented via a *singularity* container to streamline and improve the user experience, circumventing the difficulty of setting up the various software environments and dependencies.

Non-productive contigs, which are contigs that cannot be translated into a functional protein, are often filtered out by other scVDJ-seq analysis pipelines e.g. *scirpy*^13^, *scRepertoire*^15^, *Platypus*^18^ (**Supplementary Fig. 1**). In the *Immcantation/changeo*^23^ workflow, non-productive contigs are preserved and there are specific instructions for filtering or retention during annotation and clone definition steps. Moreover, *igblastn* is a V gene annotation tool^25^ and would filter contigs without V gene presence. We found that a significant proportion of contigs were non-productive in αβTCR, γδTCR and BCR data from fetal human tissues^3^ and the majority were due to absent V genes, with the exception of the TRA locus where most non-productive contigs were annotated due to presence of premature stop codons (**Fig. 2a**).

**Fig. 2.**
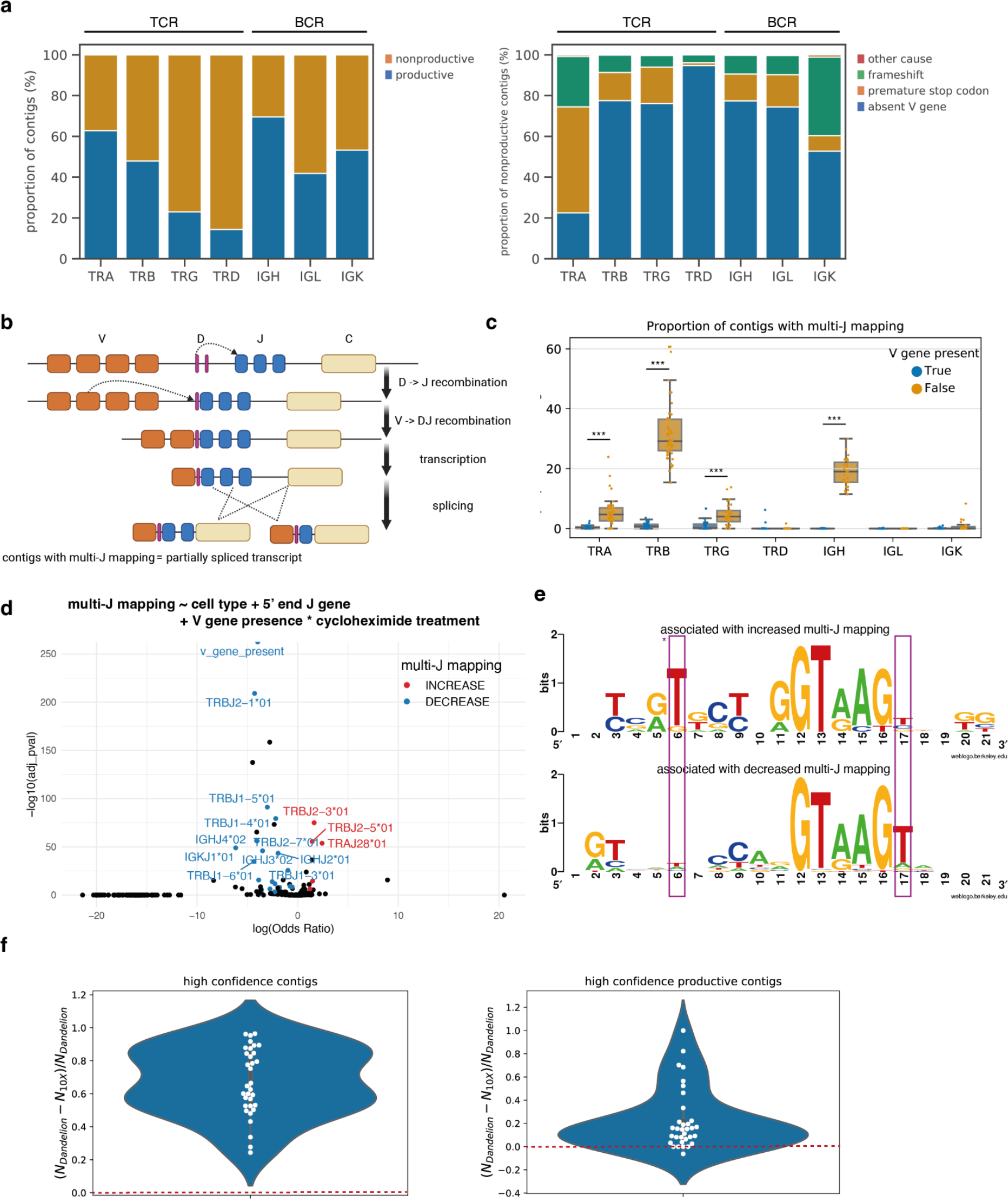
***Dandelion* offers improved contig annotations. a,** Left: barplot of proportion of contigs that are productive or non-productive in each locus. Right: barplot showing the causes of non-productive contigs in each locus. For both plots, sc-γδTCR, -αβTCR and -BCR data were taken from Suo et al. 2022^3^. **b,** Schematic illustration of the V(D)J rearrangement process and the potential cause of multi-J mapping with sequential mapped J genes on the same contig. **c,** Boxplot of the proportion of contigs with multi-J mapping, in the presence (blue) or absence (orange) of V genes. Each point represents a sample and data were taken from Suo et al. 2022^3^. Only samples with at least 10 contigs are shown. Boxes capture the first to third quartiles and whisks span a further 1.5X interquartile range on each side of the box. For each locus, the proportions in contigs with and without V genes were compared by the Wilcoxon rank sum test. *P*-values less than 0.001 were marked with *** (*P*-value for TRA: 8.78e-10; TRB: 4.96e-19; TRG: 1.47e-5; TRD: 0.68; IGH: 3.60e-11; IGL: 0.84; IGK: 0.073). **d,** Top: logistic regression formula to explore factors associated with multi-J mapping. Bottom: volcano plot summarizing logistic regression results using data from Suo et al. 2022^3^. The *y*-axis is the −log10(BH adjusted *P*-value) and the *x*-axis is log(odds ratio). The variables that were also significant in our control/cycloheximide-treated PBMC dataset were highlighted in red (associated with increased multi-J mapping) or blue (associated with decreased multi-J mapping). **e,** Sequence logos of sequences covering the last 11 nucleotides at 3′ ends (position 1 to 11) and the first 10 nucleotides of the neighboring intron (position 12 to 21) for genes associated with increased (top) or decreased (bottom) multi-J mapping. J genes associated with increased multi-J mapping were less likely to have T in position 17 (*P*-value 0.059 in logistic regression) and ‘GTAAGT’ is a known consensus motif for splicing in position 12 to 17 i.e. +1 to +6 in the intron. They were also more likely to have T in position 6 (*P*-value 0.022 in logistic regression) although the effect on splicing is unknown. **f,** Swarmplots of fraction difference of sc-γδTCR contigs annotated by *Dandelion* versus 10X *cellranger vdj* (v6.1.2) using data from Suo et al. 2022^3^. Each dot represents a sample. The red dashed line marks the threshold of 0, above which *Dandelion* recovers more γδTCR contigs than 10X. Left: all high confidence contigs. Right: high confidence productive contigs.

This pattern was consistent even after excluding thymic samples to remove the influence of developing T cells (**Supplementary Fig. 2a**). These non-productive contigs without V genes were captured in scVDJ-seq because the rapid amplification of 5′ complementary DNA (cDNA) ends (5′ RACE) technology used in the protocol does not require primers against V genes for targeted enrichment, in contrast to the previous multiplex PCR approach (**Supplementary Fig. 2b**). Although these contigs are not translated into functional proteins, they likely represent products of partial or failed recombination that we reasoned are still biologically meaningful, reflecting a cell’s history and origin. The *Immcantation* workflow would divert these contigs into a “*failed*” file and this file is not typically exposed to the user unless specified explicitly. Therefore, *Dandelion* does not automatically filter out non-productive contigs, and this data has utility, as later discussed, when we used it to track B1 cell origin and ILC/NK development.

We have also discovered that multiple J genes can be sequentially mapped onto different regions in the same messenger RNA (mRNA) contig, a phenomenon we termed ‘multi-J mapping’. Looking at the most frequent multi-J mapping contigs in each locus (**Supplementary Table 1**), we found that the majority were two to four neighboring J genes on the genome interspersed with introns. As the process of linking the chosen J to C genes is achieved through RNA splicing rather than DNA recombination, contigs with multi-J mapping are likely products of partially spliced transcripts (**Fig. 2c**). Nevertheless, it is biologically plausible that the J gene nearest to the 5′ end is the intended exon that would be expressed in the mature mRNA.

We next investigated factors that might contribute to multi-J mapping. We first noted that non-productive contigs without V genes appeared to be more likely to have multi-J mapping (**Fig. 2c**). This difference could be due to nonsense-mediated decay (NMD), an RNA degradation process that is triggered when translation encounters a premature stop codon^27^. Multi-J mapping contigs that contain a V gene will initiate translation from the V gene, which will trigger degradation by NMD due to premature stop codons in J gene introns. Transcripts of multi-J mapping without a V gene cannot be translated and will therefore evade degradation by NMD. To test the contribution of NMD to multi-J mapping, we treated peripheral blood mononuclear cells (PBMCs) with cycloheximide to block NMD and analyzed treated and untreated cells by scRNA-seq with scVDJ-seq. This resulted in an increase in the proportion of multi-J mapping in TCR contigs with V genes (**Supplementary Fig. 2c**), supporting the conclusion that NMD recognises and degrades V-gene containing multi-J mapping contigs.

We used a logistic regression model to look for additional factors associated with multi-J mapping (**Fig. 2d**) in both the Suo et al. 2022^3^ dataset (**Supplementary Table 2**) and the new control/cycloheximide-treated PBMC dataset that we generated for this study (**Supplementary Table 3**). The above finding was further supported by a significant interaction (Benjamini–Hochberg (BH) adjusted *P*-value 7.07e-04) between V gene presence and cycloheximide treatment, although the significant non-interacting V gene term (BH adjusted *P*-value 5.73e-182) in the regression fit suggests that NMD may only partially account for the effect of V genes on multi-J mapping. Furthermore, we compared the sequences of 5′ end J genes positively and negatively associated with multi-J mapping and found the known consensus motif for splicing, ‘GTAAGT’ in +1 to +6 position of adjacent intron^28^, was disrupted in J genes associated with more multi-J mapping (**Fig. 2e**, **Supplementary Table 4**). In conclusion, the factors that might contribute to multi-J mapping include specific cell types and J gene identity, which potentially affect splicing efficiencies; as well as V gene presence, which might be partially explained by NMD (illustrated by **Supplementary Fig. 2d**).

An additional application of *Dandelion*’s contig annotation functionality is that it allows for γδTCR contig annotation. There are two existing methods for sc-γδTCR mapping: i) the *cellranger vdj* pipeline developed by 10X Genomics, although this is primarily tailored for αβTCR contigs; ii) the *TRUST4*^29^ software which performs *de novo* contig assembly and annotation. The *cellranger* software is capable of reconstructing the γδTCR contigs, but most versions struggle with annotating them, a problem 10X was aware of and addressed with user-side workaround instructions (see **Supplementary Note 1**). While *TRUST4 can yield* sc- TCR annotations, including γδTCR, it relies on the presence of a V gene in the contig thus unable to handle non-productive contigs without V genes. Supplying the reconstructed contigs into *Dandelion*’s pre-processing pipeline from the cellranger output yields re-annotated output that can be used for downstream analysis. We processed 33 γδTCR libraries^3^; One mapping was done with *cellranger* 6.1.2 to the 10X GRCh38 5.0.0 V(D)J reference, with the contigs identified by *cellranger* as high confidence subsequently re-annotated with *Dandelion*. Another mapping was done with *cellranger* 6.1.2 to the 5.0.0 reference modified to obtain annotated γδTCR contigs as per 10X Genomics’ workaround instructions. We see a consistent higher recovery rate of both high confidence γδTCR contigs and high confidence productive γδTCR contigs in the mapping post-processed with *Dandelion*, verified as statistically significant by the Wilcoxon signed-rank test (*P*-value for high confidence contigs: 5.39e-7, *P*-value for high confidence productive contigs: 3.14e-6) and showing a large effect size (rank correlations equal to 1 and 0.98 for all high confidence contigs and high confidence productive contigs respectively) (**Fig. 2f**). While 10X Genomics has introduced some γδTCR support with *cellranger* 7.0.0, the results were inferior to the prior workaround from version 6 (**Supplementary Fig. 2e**).

### Creating a V(D)J feature space

To better leverage the combined gene expression and AIR repertoire data, we introduced a novel analysis strategy to create a pseudo-bulk V(D)J feature space, which transforms V(D)J data from categorical to continuous format for downstream applications (**Fig. 3a**). Transcriptionally similar cells are first grouped into pseudo-bulks, which can be based on metadata features, or partially overlapping cell neighborhoods^30^. For instance, cells can be pseudobulked by annotated cell type, donor and organ to perform differential gene usage analysis across cell types while appropriately controlling for donor and organ differences. For trajectory analysis, we recommend pseudo-bulking cells by partially overlapping cell neighborhoods sampled from gene expression space e.g. using *Milo*^30^ to model a more continuous cell state. For each pseudo-bulk, we compute the fraction of cells using each of the genes within the same segment (e.g. TRAJ1 to TRAJ61 in the TRAJ segment). These vectors of fractions are concatenated across V(D)J segments of interest to form the feature vector in the V(D)J space. This can then be used with conventional dimension reduction techniques such as principal component analysis (PCA) or uniform manifold approximation and projection (UMAP).

**Fig. 3.**
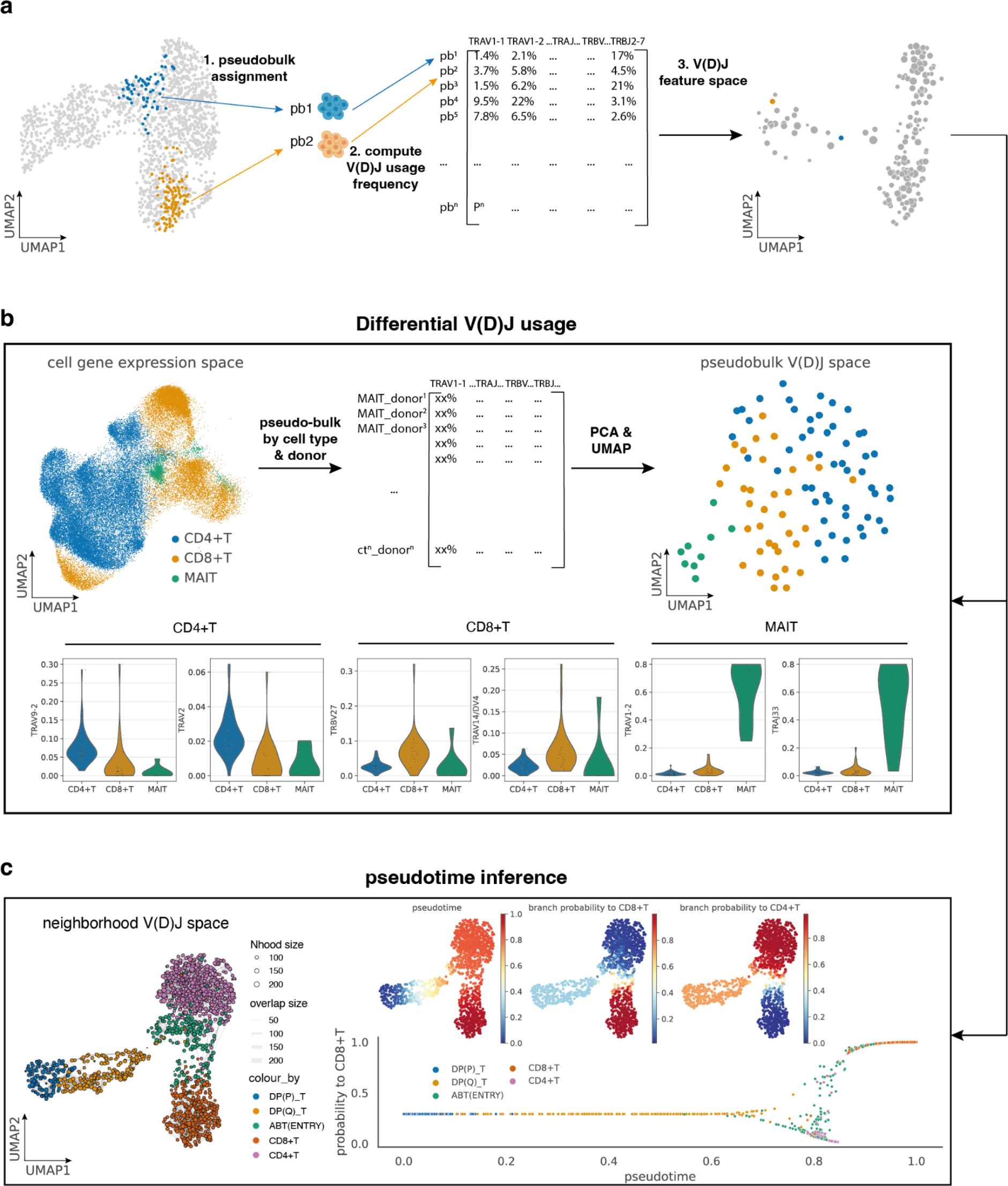
Creating a V(D)J feature space. a, Schematic illustration of the workflow of creating a V(D)J feature space. Step 1: cells are assigned to pseudo-bulks, which can be based on metadata features, or partially overlapping cell neighborhoods. Step 2: V(D)J usage frequency per pseudo-bulk is computed for each gene, and used as input of the V(D)J feature space. Step 3: the V(D)J feature space can be visualized with conventional dimension reduction techniques such as PCA or UMAP, and it can then be utilized for differential V(D)J usage analysis and pseudotime inference. **b,** Top left: gene expression UMAP of all T cells from adult human tissues in Conde et al. 2022^5^, colored by low-level cell type annotations. Each point represents a cell. Top middle: V(D)J usage frequency per celltype_donor pseudo- bulk is computed for each gene, and used as input of the V(D)J feature space. Top right: UMAP of the pseudo-bulk V(D)J feature space of the same cells. Each point represents a cell pseudo-bulk. Bottom panel: top two differentially expressed TCR genes in CD4+T cells, CD8+T cells and MAIT cells. **c,** Left: UMAP of neighborhood V(D)J feature space covering DP to mature T cells with paired productive αβTCR in data from Suo et al. 2022^3^. Each point represents a cell neighborhood, colored by the dominant cell type in each neighborhood. The point size represents neighborhood size, with connecting edges representing overlapping cell numbers between any two neighborhoods. Only edges with more than 30 overlapping cells are shown. Right top: inferred pseudotime, and branch probabilities to CD8+T and to CD4+T respectively overlaid onto the same UMAP embedding on the left. Right bottom: scatterplot of branch probability to CD8+T against pseudotime. Each point represents a cell neighborhood, colored by the dominant cell type in each neighborhood.

The utility of this V(D)J feature space is demonstrated on a dataset containing adult human T cells^5^ (**Fig. 3b**). We pseudo-bulked cells by cell types and donors to explore differential usage that is consistent across different donors. On the new UMAP computed from the V(D)J feature space, pseudo-bulks containing mucosal-associated invariant T (MAIT) cells formed a distinct cluster away from the others, in contrast to the single-cell gene expression space UMAP, indicating its unique V(D)J usage (**Fig. 3b**, **Supplementary Fig. 3a-b**). This is expected due to the semi-invariant nature of MAIT TCRs and illustrates the power of V(D)J feature space. Although there is no clear clustering in other cell types apart from MAIT (**Supplementary Fig. 3b**), there is a distinct separation between cell types that belong to CD4+T cells with those of CD8+T cells (**Fig. 3b**). The differential V(D)J usage for each cell type can be computed similarly to differentially expressed gene calculation e.g. with non-parametric statistical tests implemented within *scanpy*^14^ (**Fig. 3b**, **Supplementary Table 5**).

### Leveraging V(D)J usage in pseudotime trajectory inference

We also developed a novel usage for V(D)J data by performing pseudotime inference in lymphocytes with the cell neighborhood-based V(D)J feature space. Many pseudotime inference methods have been proposed to infer cell development based on transcriptomic similarity^31^. However, the current approaches remain problematic in immune cell development because the differentiation process is often interspersed with waves of proliferation, and transcriptomic convergence e.g. between NKT cells and NK cells can be misleading. Because usage of V(D)J genes in AIRs changes definitively as a result of cycles of recombination and selection during lymphocyte development, the AIR repertoire acts as a natural ‘time-keeper’ for developing T and B cells. A developing T cell’s fate towards CD8 *versus* CD4 T cells is determined by whether its TCR interacts with antigen presented on MHC class I or class II during positive selection. Therefore, it is biologically conceivable that the TCR gives more accurate predictions on the branch probability to each T cell lineage.

This is the motivation for leveraging V(D)J data in pseudotime inference. For this task, we chose to pseudo-bulk by cell neighborhoods as modeling cell states with partially overlapping cell neighborhoods has advantages over clustering into discrete groups; clusters do not always provide the appropriate resolution and might miss important transition states.

We sampled cell neighborhoods on a k-nearest neighbor (KNN) graph built with gene expression data using *Milo*^30^. An example is shown in **Supplementary Fig. 3c** and **Fig. 3c** using the dataset from Suo et al. 2022^3^ showing cells with paired productive αβTCR from double positive (DP) T cells to mature CD4+T and CD8+T. This neighborhood V(D)J feature space was the input to compute pseudotime with *palantir*^32^. It outputs pseudotime and branch probabilities (**Fig. 3c**) to each terminal state with a predefined starting point and terminal states (**Supplementary Fig. 3d**). The inferred pseudotime follows from proliferating DP (DP(P)) to quiescent DP (DP(Q)) T cells, to abT(entry) which splits into CD8+T and CD4+T lineages. Trends of TCR usage can also be visualized along the pseudotime trajectory (**Supplementary Fig. 3e**). Pseudotime and branch probabilities can then be projected back from neighborhoods to cells (**Fig. 4a**) by averaging the parameters from all neighborhoods a given cell belongs to, weighted by the inverse of the neighborhood size.

**Fig. 4.**
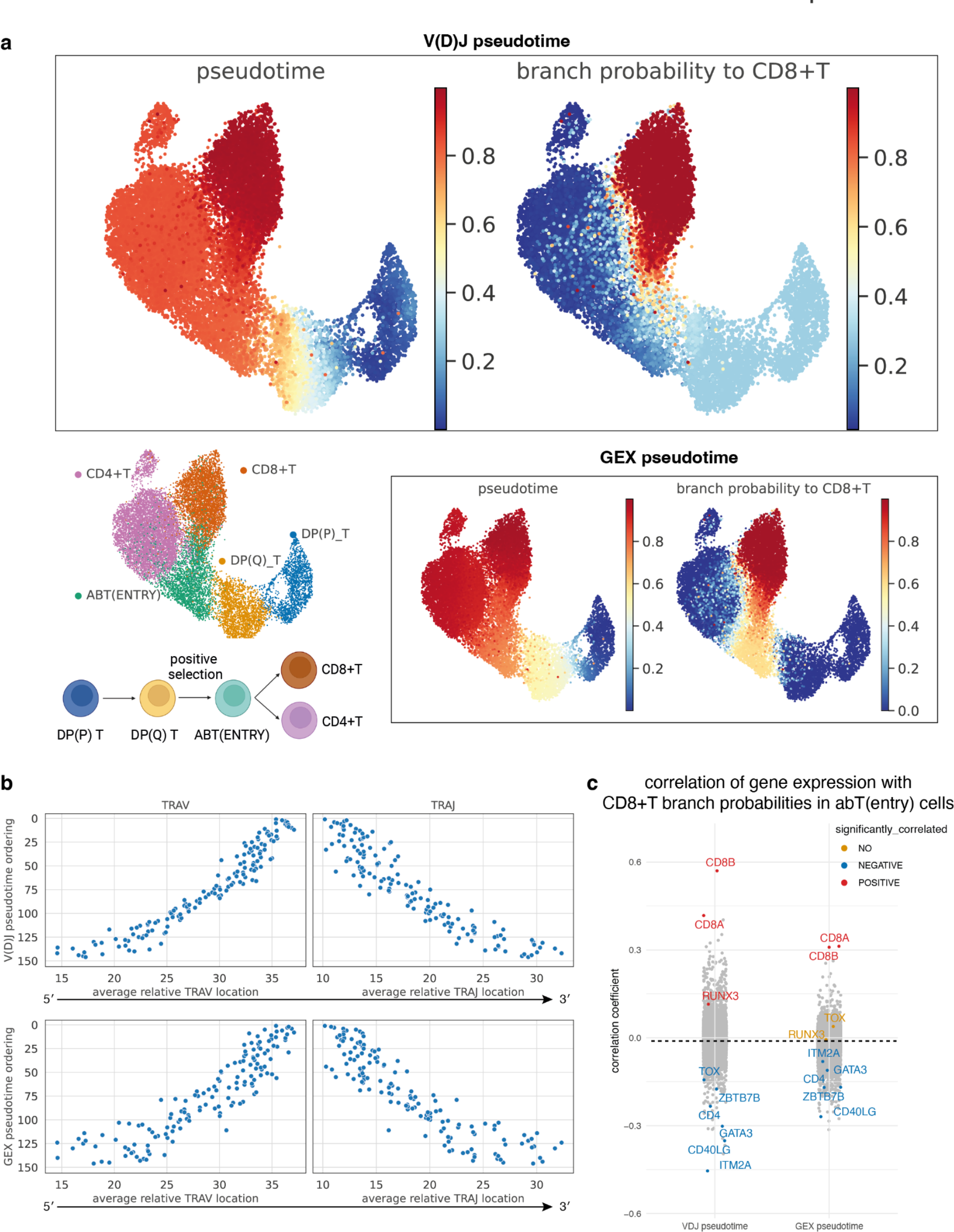
Comparing pseudotime inferred from V(D)J space or gene expression (GEX) space. a, Top: pseudotime and branch probability to CD8+T inferred from neighborhood V(D)J space in Fig. 3c, projected back to the cells, overlaid onto the same UMAP embedding as in the top left panel. Left bottom: UMAP of DP to mature T cells with paired productive αβTCR in data from Suo et al. 2022^3^. Each point represents a cell, colored by cell types. Underneath the UMAP is a schematic showing the T cell differentiation process. Right bottom: pseudotime and branch probability to CD8+T inferred from neighborhood GEX space, projected back to the cells, overlaid onto the same UMAP embedding as in the top left panel. **b,** Scatterplots of the pseudotime ordering against the average relative TRAV or TRAJ location. Each point represents a cell neighborhood. Each TRAV or TRAJ gene is encoded numerically for its relative genomic order. The *x*-axis represents the average TRAV/TRAJ relative location for each cell neighborhood. Top: results from pseudotime inferred from neighborhood V(D)J space. Bottom: results from pseudotime inferred from neighborhood GEX space. **c,** Stripplot of correlation coefficients of gene expression with branch probabilities to CD8+T within abT(entry) cells, for branch probabilities inferred from neighborhood V(D)J space and neighborhood GEX space separately. Only genes that are known CD4+/CD8+T cell markers or TFs involved in CD8+T/CD4+T lineage decision are labeled, and colored. The rest of the genes are grayed out. Labeled genes that had significant (BH adjusted *P*-value < 0.05) positive correlations were colored in red, the ones with significant negative correlations were colored in blue, and those without significant correlations were colored in orange.

There are two alternative tools, *CoNGA*^21^ and *mvTCR*^22^, that also offer integration of transcriptional profiles with TCR information. Both of them were created to detect clonally expanded cell phenotypes with CDR3 sequences being the input. We tested whether they could also be used to reveal developmental relationships with the same dataset above. With *CoNGA*, the relationships between cell types were not preserved (**Supplementary Fig. 4a**). Same applies to mvTCR (**Supplementary Fig. 4b**), which failed to separate CD4+T cells from CD8+T cells. This is not surprising, as what is changing during recombination is selection of different V(D)J genes, while CDR3 junctional sequence diversity can additionally be influenced by random nucleotide insertions. This likely explains why the sequence-based methods do not capture the intercellular relationships as faithfully as the V(D)J feature space.

### V(D)J trajectory accurately orders DP T cells and reveals early CD4/CD8 lineage decision genes

We next compared the pseudotime and branch probabilities inferred from the neighborhood V(D)J feature space with the same parameters inferred from either single-cell gene expression or neighborhood gene expression feature space.

Pseudotime inferred directly from single-cell gene expression performed unsatisfactorily, as a large proportion of CD8+T and CD4+T cells were misclassified with higher branch probabilities to the opposite terminal state (**Supplementary Fig. 5a-b**). We mainly focused our comparison with results from pseudo-bulked neighborhood gene expression (GEX) space, which produced more biologically meaningful pseudotime and branch probabilities (**Fig. 4a**). To construct the pseudo-bulked neighborhood GEX space, raw gene counts were pseudo-bulked by the same neighborhoods used to construct the V(D)J feature space (**Supplementary Fig. 3c**), and then normalized and logarithmically transformed. Pseudotime and branch probabilities were computed on this neighborhood GEX feature space and projected back to cells (**Supplementary Fig. 5c and d**). The inferred pseudotime in the pseudo-bulked space better reflected the known biology of DP(P)_T to DP(Q)_T, to abT(entry) and subsequent splits into CD8+T and CD4+T lineages. This suggests that pseudotime inference with pseudo-bulked cells work better than directly from single cells, potentially due to more stable transcriptomic profiles compared to more noisy single-cell data.

We observed two major differences when comparing the pseudotime inferred from neighborhood V(D)J feature space *versus* that from neighborhood GEX space (**Fig. 4a**). First, DP(Q) T cells appeared to dwell for a longer ‘time’ in the V(D)J trajectory as compared to the GEX trajectory. Second, the branching point of CD8+T and CD4+T cell lineages happened earlier in abT(entry) cells in the V(D)J trajectory (**Supplementary Fig. 6c**). In order to assess the fidelity of the V(D)J trajectory, we used the known fact that V-J recombination in the TRA locus happens processively^33^ using genes in the middle of the genomic locus and progressing to the two distal ends in an orderly manner. We have therefore encoded the genomic order numerically for each TRAV and TRAJ gene, and looked at the average TRAV and TRAJ relative locations for each DP(Q) neighborhood against their pesudotime ordering (**Fig. 4b**). V(D)J pseudotime showed a substantially better monotonic relationship with TRAV relative locations. Local Pearson’s correlations were computed over sliding windows of 30 adjacent neighborhoods on the pseudotime order (**Supplementary Fig. 6a**), and V(D)J pseudotime had higher absolute correlation coefficients on average (-0.67 *versus* -0.43 for TRAV). A smaller improvement was also observed for TRAJ, with the average local Pearson’s correlations improved from 0.42 to 0.50 (**Supplementary Fig. 6b**).

CD4 *versus* CD8 T cell lineage commitment is a classical immunological binary lineage decision that has been intensely investigated over many years^34^ but remains challenging to study as the selection intermediates have been difficult to observe directly^35^. We examined which genes in abT(entry) cells showed expression patterns that are correlated with branch probabilities to CD8+T lineage (**Fig. 4c**). This approach actually allows us to subdivide the abT(entry) cell population into two subsets, associated with higher probability of CD4 *versus* CD8 differentiation respectively.

When considering the top genes that were positively correlated with the CD8+ T cell lineage choice, these included *CD8A* and *CD8B*, which are markers for CD8+T cells^6^. The top genes that were negatively correlated included *CD40LG*, which is a marker for CD4+T helper cells^6^, and *ITM2A* which is found to be induced during positive selection and causes CD8 downregulation^36^. Other markers of CD4+T cells such as *CD4*^6^, together with highly validated transcription factors (TFs) that are known to be involved in CD8+T or CD4+T lineage decisions^34^, including *RUNX3*^37, 38^, *ZBTB7B*^39, 40^, *TOX*^41^ and *GATA3*^42, 43^ all displayed significant correlations in the expected directions. In contrast, when we performed the same test with CD8+T branch probabilities from GEX pseudotime, the magnitude of the correlation coefficients were notably reduced and some (e.g. *TOX* and *RUNX3*) were no longer statistically significant (**Fig. 4c**). In the case of *TOX*, the direction of the correlation was wrongly inverted (**Fig. 4c**). In addition, the V(D)J pseudotime also revealed novel associations between the trajectories and TFs such as *ZNF496*, *MBNL2* and *RORC* for CD8+T, and *SATB1, STAT5A* and *STAT1* for CD4+T (**Supplementary Fig. 6d**, full gene list in **Supplementary Table 6**). These new insights into TFs predicted to be involved in lineage commitment merit future investigations and validations.

We have also used different pseudotime inference methods to ensure the robustness of the results. Pseudotime inferred from neighborhood V(D)J space using *palantir*^32^, *monocle3*^44^, and *diffusion pseudotime*^45^ was very similar (**Supplementary Fig. 7a**) and they all showed a better monotonic relationship with TRAV or TRAJ relative locations compared to pseudotime inferred from neighborhood GEX space (**Supplementary Fig. 7b**). However, *palantir* is preferred as the only method allowing output of branch probability to each terminal state, which is useful in deciphering CD4/8 lineage decisions as shown above.

Taken together, we showed that V(D)J-based pseudotime inference gives more accurate DP(Q) T cell alignment, improves association of CD8/CD4 branch probabilities within abT(entry) cells allowing us to subdivide this cell state. We can use this approach to recapitulate known regulators, and uncover novel candidate regulators underlying CD8+T/CD4+T fate choice.

### New insights into lymphocyte development using non-productive recombination as a “fossil record”

Based on our earlier observations of high proportions of non-productive contigs being represented in the single-cell V(D)J data (**Fig. 2a**), we next explored whether different lymphoid cell types expressed different proportions of non-productive contigs. While non-productive BCR contigs were restricted to B lineage cells (**Supplementary Fig. 8a-b**) as expected, we were surprised to find that non-productive TRB contigs were not only expressed in developing DN T cells, but also in the ILC/NK lineage, and some B lineage cells (**Fig. 5a, Supplementary Fig. 8c**). The majority of the non-productive TRB contigs within ILC/NK/B cells were contigs without V gene (**Supplementary Fig. 8d**).

**Fig. 5.**
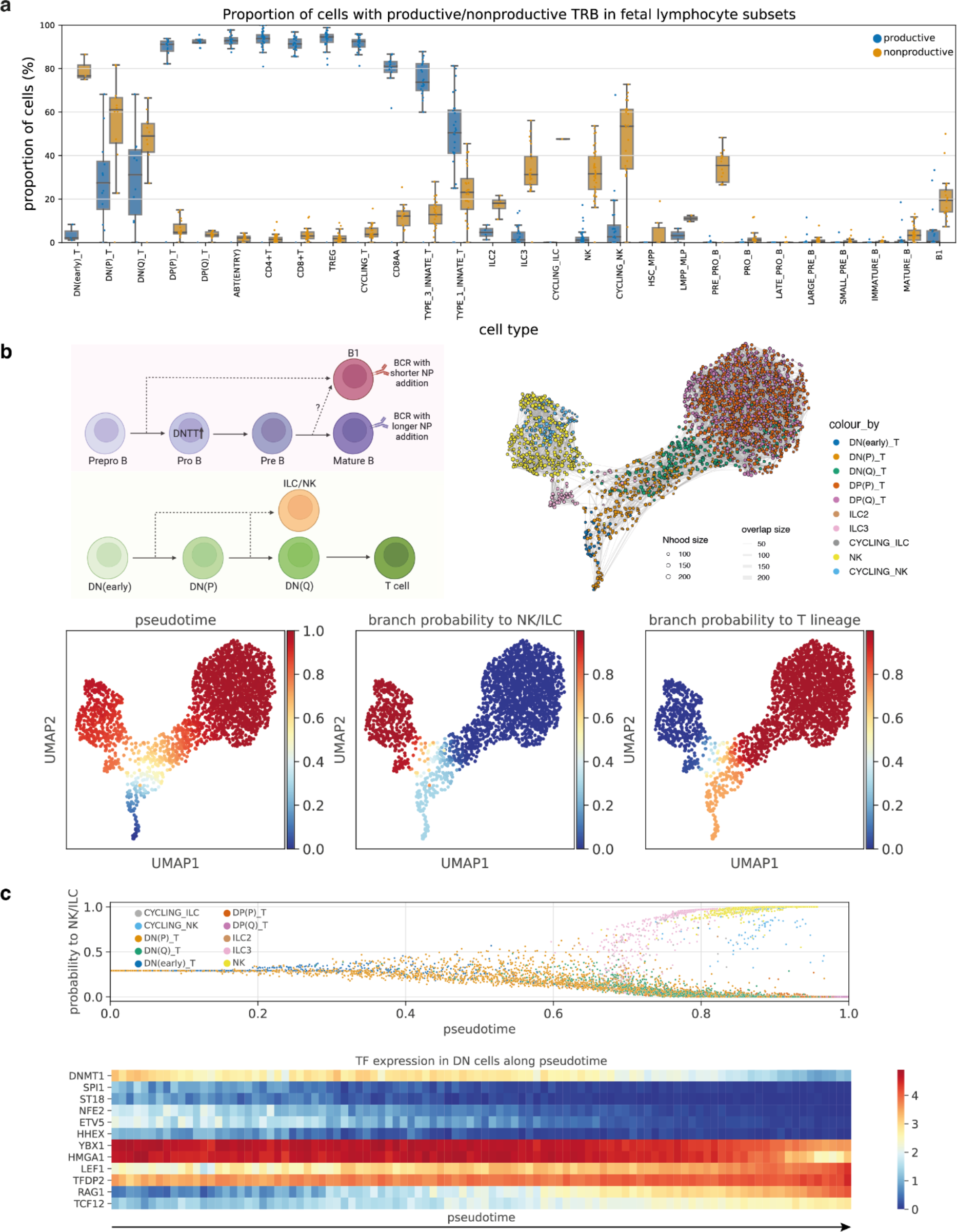
Insights into lymphocyte development from non-productive TCR. a, Boxplot of the proportion of cells with productive (blue) or non-productive (orange) TRB in different fetal lymphocyte subsets. Each point represents a sample and data were taken from Suo et al. 2022^3^. Only samples with at least 20 cells are shown. Boxes capture the first to third quartiles and whisks span a further 1.5X interquartile range on each side of the box. The annotations used here were based on the version whereby the exact identity of cycling B cells was predicted to be immature B, mature B, B1 or plasma B cells using *Celltypist*^3, 5^. The equivalent boxplot using the original annotations is shown in **Supplementary Fig. 8a**. **b,** Top left: schematic illustration showing the proposed development of B cells (top panel), and relationship between ILC/NK and T cell lineages. Top right: UMAP of neighborhood V(D)J feature space covering ILC, NK and developing T cells with TRBJ in data from Suo et al. 2022^3^. Each point represents a cell neighborhood, colored by cell types. The point size represents neighborhood size, with connecting edges representing overlapping cell numbers between any two neighborhoods. Only edges with more than 30 overlapping cells are shown. Bottom: inferred pseudotime, and branch probabilities to ILC/NK and T lineage respectively overlaid onto the same UMAP embedding on the top right. **c,** Top: scatterplot of branch probability to ILC/NK lineage against pseudotime. The pseudotime was inferred from neighborhood V(D)J space shown in Fig. 5b and projected back cells. Each point represents a cell, colored by cell types. Bottom: heatmap of TF expressions across pseudotime in DN T cells. Pseudotime is equally divided into 100 bins, and the average gene expression is calculated for DN T cells with pseudotime that falls within each bin. Genes selected here are TFs that had significantly high Chatterjee’s correlation^74^ with pseudotime (BH adjusted *P*- value < 0.05, and correlation coefficient > 0.1).

The B lineage cells with non-productive TRB contigs included pre-pro B and B1 cells but not pro- or pre-B cells (**Fig. 5a, Supplementary Fig. 8c**). Pre-pro B and B1 cells expressed only non-productive TRB but not TRG/D contigs (**Supplementary Fig. 9a-c**), suggesting that pre-pro B and B1 cells share a common development route (**Fig. 5b** schematic illustration) while bypassing pro- or pre-B cell stages. This clarifies that B1 cells in human fetal development stages can emerge through an alternative route to the rest of mature B cells (B2 cells). The conventional B cell differentiation route is thought to start from pre-pro B cells, the earliest cells that are committed to B lineage. The cells then progress through the pro- and pre-B cell stages, rearranging their BCR heavy and light chains respectively, while expressing the pre-BCR, and then emerge as immature B cells with a productive BCR and then finally differentiate into mature naive B cells^46^. Our observations are consistent with findings in murine B1s, which were shown to bypass the pre-BCR selection stage^47, 48^ that normally happens in pre-B cells to remove self-reactive B cells. This may also explain why B1 cells have BCRs with shorter non-coded/palindromic (N/P) nucleotide insertions^3^, due to negligible expression of DNTT in pre-pro B but much higher expression in pro- and late pro- B cells^3^. In addition, as pre-pro B cells are almost undetectable in adult bone marrow^49^, this potentially explains why we were unable to definitively locate B1-like cells in adult human tissues^3^.

The ILC/NK lineage also expressed non-productive TRG/D contigs with some TRA contigs (**Supplementary Fig. 9a-c**), similar to DN T cells. With the V(D)J feature space described above (**Fig. 3**), we used TRBJ frequency as the input to delineate T/ILC/NK developmental trajectories, since all of them express TRBJ (**Fig. 5b, Supplementary Fig. 10a**). The inferred trajectory suggests that ILC/NK cells deviate away from T cell development between DN(early) and DN(Q) stage (**Fig. 5b**-**c**).

Previous literature on the ILC/NK lineage has also demonstrated partial recombination of TRG/D in murine lung ILC2^50^, and of TRB/G in murine thymic ILC2^51^, leading to the hypothesis of ‘aborted’ DNs for ILC/NK development^52^. Our observation of the expression of non-productive TRB/G/D in ILC/NK cells partially supports this theory. Notably, we also observed non-productive TRB expression in ILC/NK cells in other fetal organs, with no overt differences in frequencies between organs (**Supplementary Fig. 9d**). This potentially suggests that T cells and ILC/NK cells might share the same initial stage of development, and then deviate away from each other before productive TRB/G/D is made.

In addition, by examining the expression patterns of transcription factors (**Fig. 5c**) and genes encoding cell surface proteins (**Supplementary Fig. 10b**) that changed along the TRBJ- inferred pseudotime, we can define stages for DN development at higher resolution than previously reported in the literature. We observed that expression levels of genes such as *SPI1*, *RAG1*, *HHEX*, *TCF12*, *CD34*, *CD3D*, *CD8A*, *CD8B* followed an expected pattern along the trajectory^53^. At the same time, we also discovered many novel genes that could re- define DN stages. We further noted that there were some discordances in expression patterns of selected transcription factors between human and mouse DN development^53^ (**Supplementary Fig. 10c**). However, this discrepancy could be due to age mismatch i.e. fetal human to adult mouse, and the murine data was mainly learnt from transcription factor knockout instead of scRNA-seq studies. Comparison across different human age groups, as well as a detailed comparison with mouse thymic scRNA-seq with paired scVDJ-seq data need to be performed in the future.

Finally, we have also explored BCR expression in plasmacytoid dendritic cells (pDC). Several papers have reported that pDC can be derived from both myeloid and lymphoid progenitors^54, 55^ and there is IgH D-J rearrangement in some pDCs^54, 56–59^. The question is whether the pDCs that have initiated BCR rearrangements are derived from lymphoid progenitors^54, 55^. We have repeated the analysis in all the myeloid cells in the human fetal dataset^3^. There are indeed some non-productive BCR in pDC (both heavy and light chain as shown in **Supplementary Fig. 11a and b**), in agreement with previously reported IgH D-J rearrangement in pDC^54, 56–59^. However, pDC itself expresses RAG and DNTT (**Supplementary Fig. 11c**). This means that the presence of non-productive BCR does not necessarily indicate that pDCs are derived from lymphoid progenitors as BCR rearrangement can be carried by RAG in pDC itself, similar to what has been claimed in Shigematsu et al. 2004^57^. While it may be interesting to use our VDJ-based trajectory to explore whether the development of pDC overlaps with early B cell development, the current dataset is limited by the cell number as only 51 pDC and cycling pDC cells have non-productive IGH. This is certainly worth future investigations when the data becomes available.

In summary, the unexpected finding of expression of non-productive TCR contigs in specific cell types has the potential to shed new light on the origin and history of lymphocyte development. We have utilized this information and suggested that B1 potentially arises directly from pre-pro B cells, and provided support for the ‘aborted’ DN theory for the origin of ILC/NK cells.

## Discussion

Overall, *Dandelion* improves upon existing methods with more refined contig annotations, recognising non-productive contigs, identifying multi-J mapping and recovering more γδTCR contigs. In conjunction with our novel V(D)J feature space approach with pseudotime trajectory inference, it has allowed us to better align CD4 *versus* CD8 T cell lineage commitment processes, and further identify developmental origins of innate-like lymphocyte cells.

Our improved data processing workflow revealed two unexpected data challenges and opportunities with scVDJ-seq. First, the surprising observation that a high proportion of TCR/BCR contigs are non-productive suggests that these are unique data challenges in the single-cell space due to choice of library construction. However, it is not unexpected as V(D)J rearrangement is a ‘wasteful’ exercise, a price that comes with the generation of effective and diverse immune response; for example, two out of three rearrangement events for immunoglobulins are destined to be non-productive^60, 61^. While non-productive TCRs and BCRs from high-throughput ‘bulk’ AIR sequencing data have previously been used in conjunction with productive contigs to estimate the generation probabilities and diversities of AIRs during affinity maturation and infection^62, 63^, these would only have factored in those with V gene annotation due to library construction limitations. Through scVDJ-seq and analysis using *Dandelion*, we now have the ability to corroborate this at the single-cell level, including partially rearranged contigs, as outlined in our analysis of innate lymphocyte development. This suggests that the presence of the non-productive contigs may have important biological implications in a cell-type specific manner.

Second, detection of multi-J mapping suggests that these are naturally occurring and likely represent products of partial splicing events at the transcript level. A few factors were identified to be associated with multi-J mapping, including J gene identities, which potentially affect splicing efficiencies with their disrupted splicing site, as well as V gene presence, which might be partially explained by NMD^27^. The biological implications of the presence of these multi-J mapping contigs are unclear at this stage and require future experimental validation to understand how and why they arise.

We introduced a novel way of analyzing the single-cell V(D)J modality in *Dandelion* with the pseudo-bulk V(D)J feature space, which can be used for visualization and differential V(D)J usage testing. In addition, when the pseudo-bulking is done by gene expression neighborhoods, the V(D)J feature space is anchored to the underlying gene expression feature space where cell neighborhoods are sampled. We utilized this approach for pseudotime trajectory inference and demonstrated its advantages in both of our case studies.

The first case study examined the processes underlying T cell development in the thymus. Our approach allowed us to discover that fate commitment starts earlier than expected with the inclusion of TCR information. It was previously suggested that abT(entry) cells were likely to be a point of divergence due to its position as an intermediary cell state between DP T cells and mature single positive T cells^6^. With this new technique that includes TCR information, we are now able to better delineate the branching point to a much earlier point within the abT(entry) cells. The gene expression patterns of marker genes and transcription factors known to be associated with CD4 *versus* CD8 T cell fate were better aligned with the new trajectories. Our analysis has further revealed novel CD4/8 associations with other transcription factors that remain to be explored.

Similar approaches can be applied to other TCR trajectories in different contexts e.g. across different developmental stages in human lifespan, diseases and *in vitro* settings. It remains to be seen whether a VDJ-based trajectory can be utilized in T cell activation. Furthermore, this approach has not been optimized for BCR trajectories, as we are limited by the small number of B progenitors in the existing dataset collections. Further, BCRs have additional rearrangement rules that need to be considered e.g. somatic hypermutation, differential rearrangement events leading to asymmetric usage of kappa and lambda light chains and light chain editing processes^64^, as well as recently described light chain coherence in functional antibodies^65^. We hope to improve on these aspects in a future iteration of *Dandelion* when more single-cell V(D)J data become available.

The second case study extended the observations of non-productive V(D)J contig representation in 10X Genomics’ single-cell data, which has been largely ignored and/or not easily accessible with other existing workflows e.g. *scirpy*^13^ and *immcantation*^23^. Our unexpected finding that B1 cells and pre-pro B cells were expressing relatively higher levels of non-productive TRB contigs suggest that B1 lineage commitment diverged earlier than expected, some time between the pre-pro B stage and pro-B stage. Two competing models have been described with regard to B1 origin^66^. The lineage model, or layered immune system hypothesis^67^ proposed that B1 cells compared to B2 cells arise from distinct progenitors emerging at different times during development^68–71^; while the selection model hypothesized that they originate from the same progenitors but after differential signaling depending on self-reactivity^72, 73^. Our finding here potentially offers a reconciliation of both models, with fetal specific pre-pro B cells being B1 progenitors supporting the layered immune system model, and the skipping of pre-BCR selection presumably allows formation of self-reactive BCR in support of the selection model.

The enrichment of the non-productive TRB contigs is not just found in the pre-pro B and B1 cells, but also in NK and ILC lineage cells along with non-productive TRG and TRD. The latter lineage is easier to explain as partial recombination of TCR has been reported in murine ILC^50, 51^ and our findings support the ‘abandoned’ DN theory^52^. The hypothesis is that ILC/NK cells are originally on a canonical T cell development trajectory, but subsequently influenced to abort this process, resulting in sustained expression of non-productive TCR rearrangements whilst developing into ILC/NK. Perhaps this is driven by overexpression of key transcription factors such as *ID2* and *ZBTB16*^52, 53^, or lack of NOTCH signaling^52^. While we cannot rule out other routes of ILC/NK development, our new insights do support the notion that T and NK/ILC developments partially overlap but diverge before productive TCRs are rearranged. Our analysis offers new insights into transcription factors and surface marker genes that define DN T cell stages at high resolution, opening avenues for future in-depth investigation.

In summary, we present *Dandelion* as an easy-to-use package/pipeline for integrative analyses of single-cell GEX and V(D)J data modality. The package is freely available online at https://github.com/zktuong/dandelion with tutorials and demo cases and is actively updated for further improvements. The pseudo-bulk V(D)J data is also publicly available for use as a reference to project or align new query data e.g. for disease samples such as cancers that originate from T cells. We hope that the software and the resource will be useful to the community for exploring lymphocyte biology in the single-cell space, generating new insights that will help advance our understanding of immune cell development and function in health and disease.

## Methods

### Dandelion

#### Pre-processing

*Dandelion* can run the pre-processing of data using the standard outputs from all *cellranger vdj* versions. In this manuscript, single-cell V(D)J data from the 5′ Chromium 10X kit were initially processed with *cellranger vdj* pipeline (v6.1.2) with *cellranger vdj* reference (v5.0.0). TCR and BCR contigs contained in ‘*all_contigs.fasta*’ and ‘*all_contig_annotations.csv*’ from all three library types (αβTCR, γδTCR and BCR) were then reannotated using an *immcantation*-inspired^23^ pre-processing pipeline contained in the *Dandelion* singularity container (v0.3.0).

The pre-processing pipeline includes the following steps:

i. adjust cell and contig barcodes by adding user-supplied suffixes and/or prefixes to ensure that there are no overlapping barcodes between samples;
ii. optionally subset to contigs deemed high confidence in the *cellranger* output; this was done in the analysis performed here;
iii. re-annotation of contigs with *igblastn* (v1.19.0) against IMGT (international ImMunoGeneTics) reference sequences (last downloaded: 01/08/2021) with the following parameters: minimum D gene nucleotide match = 9, V gene e-value cutoff = 10^-4^; rearrangements missing the CDR3/junction sequences are enforced to be non-productive (productive = “F”) and incomplete (complete_vdj = “F”).
iv. re-annotation of D and J genes separately using *blastn* with similar parameters as per *igblastn*^25^ (dust =“no”, word size (J = 7; D = 9)) but with an additional e-value cutoff (J = 10^-4^ in contrast to *igblastn*’s default cut off of 10; D = 10^-3^). This is to enable annotation of contigs without the V gene present;
v. identification and recovery of non-overlapping individual J gene segments (under associated ‘*j_chain_multimapper*’ columns). In the list of all mapped J genes (*all_contig_j_blast.tsv*) from *blastn*, the J gene with the highest score (*j_support*) was chosen. *Dandelion* then looks for the next J gene with the highest ‘*j_support*’ value, and with start (*j_sequence_start*) and end (*j_sequence_end*) position not overlapping with the selected J gene, and does so iteratively until the list of all mapped J genes is exhausted. In contigs without V gene annotations, we then select the 5′ end leftmost J gene and update the ‘*j_call*’ column in the final AIRR table. For contigs with V gene annotations, but with multiple J gene calls, we use the annotations provided by *igblastn* (NCBI IgBLAST Release 1.19.0’s release notes states that they “*\*Added logic to handle the case where there is an unrearranged J gene downstream of the VDJ rearrangement.*”).

For BCRs, there are two additional steps:

vi) additional re-annotation of heavy-chain constant (C) region calls using *blastn* (v2.13.0+) against curated sequences from CH1 regions of respective isotype class;
vii) heavy chain V gene allele correction using tigger (v1.0.0)^75^. The final outputs are then parsed into AIRR format with *change-o* scripts^23^.

All the outputs from each step are saved in a subfolder which the user can elect to retain or remove as per their requirements. Typically a user would proceed with the file ending with the suffix ‘*_contig_dandelion.tsv*’ as this represents the rearrangement sequences that pass standard quality control checks. In this manuscript, we used the data found in the ‘*all_contig_db-all.tsv*’ as it also contains the multi-J mapping.

#### Post-processing

In addition to the pre-processing steps at the contig level, post-processing, or integrating cell-level quality control, is performed using *Dandelion*’s ‘*check_contig*’ function. The function checks through whether a rearrangement is annotated with consistent V, D, J and C gene calls and performs special operations when a cell has multiple contigs. All contigs in a cell are sorted according to the unique molecular identifier (UMI) count in a descending order and productive contigs are ordered higher than non-productive contigs. For cells with other than one pair of productive contigs (one VDJ and one VJ), the function will assess if the cell is to be flagged with having orphan (no paired VDJ or VJ chain), extra pair(s) or ambiguous (biologically irreconcilable e.g. both TCRs and BCRs in the same cell) status with some exceptions: ii) IgM and IgD are allowed to co-exist in the same B cell if no other isotypes are detected; ii) TRD and TRB contigs are allowed in the same cell because rearrangement of TRB and TRD loci happens at the same time during development and TRD variable region genes exhibits allelic inclusion^76^. The function also asserts a library type restriction with the rationale that the choice of the library type should mean that the primers used would most likely amplify only relevant sequences to a particular loci. Therefore, if there are any annotations to unexpected loci, these contigs likely represent artifacts and will be filtered away. A more stringent version of ‘*check_contigs*’ is implemented in a separate function, ‘*filter_contigs*’, which only considers productive VDJ contigs, asserts a single-cell should only have one VDJ and one VJ pair, or only an orphan VDJ chain, and explicitly removes contigs that fail these checks (with the same exceptions for IgM/IgD and TRB/TRD as per above). If a single-cell gene expression object (*AnnData*) is provided to the functions, it will also remove contigs that do not match to any cell barcodes in the gene expression data.

Lastly, *Dandelion* can accept any AIRR-formatted data formats e.g. BDRhapsody VDJ data.

#### Clonotype definition and diversity

*Dandelion*’s mode of clonotype definition and network based diversity analysis has been previously described^4^. Briefly, TCRs and BCRs are grouped into clones/clonotypes based on the following sequential criteria that apply to both heavy-chain and light-chain contigs: (1) identical V and J gene usage; (2) identical junctional CDR3 amino acid length; (3) CDR3 sequence similarity: for TCRs, 100% nucleotide sequence identity at the CDR3 junction is recommended while the default setting for BCRs is to use 85% amino acid sequence similarity (based on Hamming distance). Single-cell V(D)J networks are constructed using adjacency matrices computed from pairwise Levenshtein distance of the full amino acid sequence alignment for TCR/BCR(s) on a per cell basis. A minimum-spanning tree is then constructed on the adjacency matrix for each clone/clonotype, creating a simple graph with edges indicating the shortest total edit distance between a cell and its neighbor. Cells with total pairwise edit distance of zero are then connected to the graph to recover edges trimmed off during the minimum-spanning-tree construction step. A graph layout is then computed either using the Fruchterman–Reingold algorithm in *networkx* (≥ v2.5) or Scalable Force-Directed Placement algorithm implemented through *graph-tool* package^77, 78^. Visualization of the resulting single-cell V(D)J network is achieved via transfer of the graph to relevant ‘*AnnData*’ slots, allowing for access to plotting tools in *scanpy*. The resulting V(D)J network enables computation of Gini coefficients based on cluster/cell size/centrality distributions, as discussed previously^4^.

### Pseudo-bulk V(D)J feature space

Pseudo-bulk construction requires pseudo-bulk assignment information of cells, along with V and J genes for the cells’ identified primary TCR/BCR contigs (selected based on productive status and highest UMI count). The former is a cell by pseudo-bulk binary matrix which can be either explicitly provided by the user or inferred from unique combinations of cell level discrete metadata. While the code is calibrated to work with *Dandelion*’s structuring by default, it can work with any V(D)J processing provided it stores cell level information on primary per-locus V/D/J calls. The input is used to generate a pseudo-bulk by V(D)J feature space, with the V(D)J calls converted to a binary matrix, added up for each pseudo-bulk, and normalized to a unit sum on a per-pseudo-bulk, per-locus, per-segment basis. The cell by pseudo-bulk information is stored in the resulting object for potential communication with the original cell space. Utility functions are provided for compatibility with *palantir*^32^ output for trajectory inference.

### Non-productive TCR/BCR contigs

Single-cell BCR, αβTCR and γδTCR data from Suo et al. 2022^3^ were remapped with *cellranger vdj* (v6.1.2) and processed further using *Dandelion* as described above. For all samples, contigs were extracted from ‘*all_contig_igblast_db-all.tsv*’ or in the case whereby ‘*all_contig_igblast_db-all.tsv*’ was empty, ‘*all_contig_igblast_db-fail.tsv*’ was used.

Preprocessed and annotated scRNA-seq data was downloaded from https://developmental.cellatlas.io/fetal-immune. Only contigs from annotated cells were kept for downstream analysis. For each contig, productive status was obtained from the column ‘*productive*’, and the causes for non-productive contigs were extracted from ‘*vj_in_frame*’ (is ‘F’ if there is a frameshift), ‘*stop_codon*’ (is ‘T’ if there is a premature stop codon) and ‘*v_gene_present*’ (is ‘False’ if V gene is absent) columns.

### Cycloheximide treatment on PBMC

A vial of frozen PBMCs was acquired from Stemcell Technologies with informed consent (as stated by Stemcell Technologies) and approval from the Yorkshire & The Humber - Leeds East Research Ethics Committee (19/YH/0441). Frozen PBMCs were thawed in pre-warmed RF10 media, which was RPMI (Sigma-Aldrich) supplemented with 10% fetal bovine serum (FBS; Gibco) and penicillin/streptomycin (Sigma-Aldrich). Cells were pelleted by centrifugation at 500g for 5 min and resuspended in RF10 media, and split between two 10 cm petri dishes. Control PBMCs were then incubated in a total of 10 ml RF10 media at 37°C for 2 hr, whereas treated PBMCs were incubated in RF10 supplemented with cycloheximide

(Sigma-Aldrich; final concentration of 100 μg/ml). After incubation, control and treated PBMCs were washed with ice cold RF10 and resuspended in 2% FBS in phosphate buffered saline (PBS; Gibco). For treated PBMCs, both the washing and resuspension buffer contained 100 μg/ml cycloheximide.

Control and treated PBMCs were then loaded onto two separate channels of the Chromium chip from Chromium single cell V(D)J kit (10X Genomics 5′ v2) following the manufacturer’s instructions before droplet encapsulation on the Chromium controller. Single-cell cDNA synthesis, amplification, gene expression (GEX) and targeted BCR and αβTCR libraries were generated. Sequencing was performed on the Illumina Novaseq 6000 system. The gene expression libraries were sequenced at a target depth of 50,000 reads per cell using the following parameters: Read1: 26 cycles, i7: 8 cycles, i5: 0 cycles; Read2: 91 cycles to generate 75-bp paired-end reads. BCR and TCR libraries were sequenced at a target depth of 5000 reads per cell.

Raw scRNA-seq reads were mapped with *cellranger* 3.0.2 with Ensembl 93 based GRCh38 reference. Low quality cells were filtered out (minimum number of reads > 2000, minimum number of genes > 500, maximum number of genes < 7000, maximum mitochondrial reads fraction < 0.2, maximum *scrublet*^79^ (v0.2.1) doublet score ≤ 0.5). Data normalization and log transformation were performed using *scanpy*^14^ (v1.9.1) (*scanpy.pp.normalize_per_cell(counts_per_cell_after=10e4*) and *scanpy.pp.log1p*). Highly variable genes were then selected (*scanpy.pp.highly_variable_genes*), and PCA (*scanpy.pp.pca*), neighborhood graph (*scanpy.pp.neighbors*) and UMAP (*scanpy.tl.umap*) were computed. Automatic annotation was done using *celltypist* (v1.2.0) (*celltypist.annotate(model = ’Immune_All_Low.pkl’, majority_voting = True)*).

Single-cell αβTCR and BCR sequencing data was mapped with *cellranger vdj* (v6.1.2) and processed further using *Dandelion* as described above. For all samples, contigs were extracted from ‘*all_contig_igblast_db-all.tsv*’ or in the case whereby ‘*all_contig_igblast_db- all.tsv*’ was empty, ‘*all_contig_igblast_db-fail.tsv*’ was used. Only contigs from annotated cells were kept for downstream analysis.

### Factors associated with multi-J mapping

#### Logistic regression analysis

We used the following logistic regression model to look for factors associated with multi-J mapping:

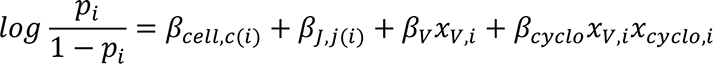

where *p*_*i*_ is the probability of multi-J mapping present in the *i*th contig, c(i) and j(i) are the cell type and the 5′ end J gene of the *i*th contig respectively, *x*_*v,i*_ is the indicator of whether V gene is present in the *i*th contig and *x*_*cyclo,i*_ is the indicator of whether *i*th contig belongs to a cell that had cycloheximide treatment. Here, (*β*_*cell, c*_: *c* ∈ *cell types*), (*β*_(j,j)_: *j* ∈ 5′ *end J genes*), *β*_*v*_ and *β*_*cyclo*_ are parameters to be estimated.

To control for multiple testing, *P*-values were adjusted with Benjamini–Hochberg procedure^80^. This was applied on all contigs from the γδTCR, αβTCR and BCR sequencing data that were identified within high-quality annotated cells from Suo et al. 2022^3^ and results are shown in **Supplementary Table 2**; and it was also applied on contigs from the αβTCR and BCR sequencing data that were identified within high-quality annotated cells from control/cycloheximide-treated PBMCs and results are shown in **Supplementary Table 3**.

#### Splicing site motif analysis

For the lists of 5′ end J genes that had significant (BH adjusted *P*-value < 0.05) association with increased or decreased multi-J mapping from **Supplementary Table 2**, the sequences of the last 11 nucleotides at each gene’s 3′ ends with the first 10 nucleotides of its 3′ end intron were extracted from the 10X GRCh38 2020-A reference. Sequence logos shown in **Fig. 2e** were generated on https://weblogo.berkeley.edu/logo.cgi81.

### γδTCR annotation comparison

To compare our γδTCR annotations against the 10X *cellranger vdj* output in the 33 γδTCR libraries^3^, we performed two additional mappings following 10X γδTCR support instructions. In one, the 5.0.0 reference was modified according to 10X instructions by replacing all instances of TRG with TRA and TRD with TRB. The reference was filtered to just TRG/TRD sequences prior to this replacement to avoid erroneous sequence overlaps. For the other, we performed the alignment with *cellranger* v7.0.0 with the accompanying reference (v7.0.0). The output of these two mappings was compared with the *cellranger* - *Dandelion* pre-processing pipeline described above. The number of high confidence γδTCR contigs and high confidence productive γδTCR contigs were determined for each mapping and each sample, and mappings were compared with the Wilcoxon signed-rank test. The effect size r is the rank correlation, which is the signed-rank test statistic divided by the total rank sum^82^.

### Differential V(D)J usage in adult T cell subsets

Preprocessed and annotated scRNA-seq data of T and innate lymphoid cells with paired αβTCR information from Conde et al. 2022^5^ was downloaded from https://www.tissueimmunecellatlas.org/. Only cells within the T cell subsets with paired αβTCR were included in the downstream analysis. T_CD4/CD8 was excluded as a low quality cell cluster. The cells were then pseudo-bulked by donor ID and cell type, and the pseudo-bulk V(D)J feature space was created with TRAV, TRAJ, TRBV and TRBJ. Only pseudo-bulks with at least 10 cells were kept. PCA, neighborhood graph and UMAP of the pseudo-bulk V(D)J feature space were computed using *scanpy*^14^ (v1.9.1) with default settings (*scanpy.pp.pca*, *scanpy.pp.neighbors*, *scanpy.tl.umap*).

For low-level cell type annotations, Tem/emra_CD8, Tnaive/CM_CD8, Trm/em_CD8, Trm_gut_CD8 were grouped into CD8+T, and Teffector/EM_CD4, Tfh, Tnaive/CM_CD4, Tnaive/CM_CD4_activated, Tregs, Trm_Th1/Th17 were grouped into CD4+T, while MAIT was left as a separate annotation. For differential V(D)J usage, Wilcoxon rank-sum test was performed using *scanpy.tl.rank_genes_groups(method=’wilcoxon’)*.

### Pseudotime inference from DP to mature T cells

#### Data integration and filtering

scRNA-seq data of human fetal lymphoid cells from Suo et al. 2022^3^ was integrated with *Dandelion* preprocessed αβTCR, BCR and γδTCR data (see section on **Non-productive TCR/BCR contigs**, using *all_contig_igblast_db-all.tsv* for all samples) with *dandelion.tl.transfer*. Two samples from F67, F67_TH_CD137_FCAImmP7851896 and F67_TH_MAIT_FCAImmP7851897 were excluded from the analysis as they were sorted for specific T cell subpopulations, instead of the CD45 sorting in all other donor samples, and inclusion might result in biased TCR sampling within this donor. Only DP(P)_T, DP(Q)_T, ABT(ENTRY), CD8+T, CD4+T cells with productive TRA and TRB were included for the trajectory analysis. Neighborhood graph (*scanpy.pp.neighbors(n_neighbors = 50)*) and UMAP (*scanpy.tl.umap*) was re-calculated using scVI latent factors as the initial data was integrated with *scVI*^83^.

#### Pseudotime inference from neighborhood V(D)J feature space

Neighborhoods were sampled using *Milo*^30^ (milopy v0.1.0) (*milo.make_nhoods*). Cells were pseudo-bulked by the sampled neighborhoods and the V(D)J feature space was created with cells’ primary TRAV, TRAJ, TRBV and TRBJ genes. The cell type annotation of each neighborhood was assigned to be the most frequent annotation of the cells within that neighborhood. PCA, neighborhood graph and UMAP of the neighborhood V(D)J feature space were computed using *scanpy*^14^ (v1.9.1) with default settings (*scanpy.pp.pca*, *scanpy.pp.neighbors*, *scanpy.tl.umap*).

For pseudotime trajectory analysis, *palantir*^32^ (v1.0.1) was used and diffusion map was computed using the first five principal components (PCs) (*palantir.utils.run_diffusion_maps(n_components=5)*, *palantir.utils.determine_multiscale_space*). The root cell was chosen to be the DP(P) T neighborhood with the smallest value on UMAP1 axis, and the two terminal states were chosen with the largest and smallest values on the UMAP2 axis for CD4+T and CD8+T neighborhoods respectively (**Supplementary Fig. 3d**). Pseudotime and branch probabilities to the terminal states were then computed with *palantir.core.run_palantir(num_waypoints=500)*.

Imputed pseudotime and branch probabilities were then projected back from neighborhoods (**Fig. 3c**) to cells (**Fig. 4a** top panel) by averaging the parameters from all neighborhoods a given cell belongs to, weighted by the inverse of the neighborhood size. Cells that did not belong to any neighborhood were removed (91 out of 17248).

For pseudotime inferred with other trajectory inference methods as shown in **Supplementary Fig. 7,** *monocle3*^44^ (0.2.3.0) was applied on the UMAP embedding of the neighborhood V(D)J feature space and *diffusion pseudotime*^45^ was applied using *scanpy.tl.dpt* function with default settings. The same root cell neighborhood was used as above.

### Pseudotime inference from neighborhood GEX feature space

Raw gene counts from scRNA-seq data were pseudo-bulked by the same cell neighborhoods as above. Data normalization and log transformation were performed using *scanpy*^14^ (v1.9.1) (*scanpy.pp.normalize_per_cell(counts_per_cell_after=10e4*) and *scanpy.pp.log1p*). Highly variable genes were then selected (*scanpy.pp.highly_variable_genes*), and PCA (*scanpy.pp.pca*), neighborhood graph (*scanpy.pp.neighbors*) and UMAP (*scanpy.tl.umap*) of the neighborhood GEX feature space were computed. Pseudotime trajectory inference was done similar to above with the first five PCs. The root cell was chosen to be the DP(P) T neighborhood with the smallest value on UMAP1 axis, and the two terminal states were chosen with the smallest and largest values on the UMAP2 axis for CD4+T and CD8+T neighborhoods respectively (**Supplementary Fig. 5c**). Imputed pseudotime and branch probabilities were then projected back from neighborhoods (**Supplementary Fig. 5d**) to cells (**Fig. 4a** bottom right panel).

### Pseudotime inference from single cell GEX

Pseudotime trajectory inference was performed with *palantir*^32^ (v1.0.1) using the first 20 scVI latent factors. The root cell was chosen to be the DP(P) T cell with the largest value on UMAP2 axis, and the two terminal states were chosen with the largest value on the UMAP2 axis for CD8+T and smallest value on the UMAP1 axis for CD4+T cells respectively (**Supplementary Fig. 5a**). Results of the inferred pseudotime and branch probabilities are shown in **Supplementary Fig. 5b**.

### Correlation between pseudotime ordering and relative TRAV/TRAJ locations

The relative genomic location of each TRAV gene was encoded numerically based on its order among all TRAV genes from 5′ to 3′ on the genome, and similarly for TRAJ. For each neighborhood, its relative TRAV or TRAJ location was computed by the average relative locations of all cells within that neighborhood. Only neighborhoods that had more than 90% cells being DP(Q) T cells were selected. The relative pseudotime order was plotted against the average relative TRAV or TRAJ location for each neighborhood in **Fig. 4b**. Local Pearson’s correlations were then computed over sliding windows of 30 adjacent neighborhoods on the pseudotime order (**Supplementary Fig. 6a-b**).

*<u>Correlation between gene expression and branch probabilities to CD8+T in abT(entry) cells</u>* Pearson’s correlations were computed between gene expression and branch probabilities to CD8+T lineage within abT(entry) cells for all genes. *P*-values were adjusted for multiple testing with Benjamini–Hochberg procedure. Results are shown in **Fig. 4c**, **Supplementary Fig. 6d** and **Supplementary Table 6**.

### VDJ-based dimensionality reduction with *CoNGA*^21^

Preprocessed and annotated scRNA-seq data of human fetal lymphoid cells from Suo et al. 2022^3^ was downloaded from https://developmental.cellatlas.io/fetal-immune. Matching αβTCR samples had their *all_contig_annotations.csv cellranger* output files flagged with the sample IDs for both cell and contig IDs, and were subsequently merged into a single file and subset to just high confidence contigs for cells present in the scRNA-seq object. This file was used on input for *CoNGA*’s (v0.1.1) setup_10x_for_conga.py script, which produced a tcrdist-based PCA representation of the cells’ VDJ data. The PCA coordinates were used to compute a neighborhood graph and UMAP representation (**Supplementary Fig. 4a**), using default *scanpy* settings.

### Joint embedding of single cell gene expression and TCR with *mvTCR*^22^

The same cells for which we performed pseudotime inference from DP to mature T cells above were used in the *mvTCR* (version under development, cloned from the repo at commit 528d3e11a360fc4b0f09d782b88f5ec7de9283d6) trial. Clonotypes were called based on CDR3 nucleotide sequence identity of the cells’ primary TRA and TRB chains (*scirpy.pp.ir_dist*, and *scirpy.tl.define_clonotypes(receptor_arms=”all”*, *dual_ir=”primary_only”)*).

Normalized and log transformed data was used as recommended in *mvTCR*’s tutorial. The donor ID was one-hot encoded and supplied as a conditional variable. 80% of cells were used as training data, the remaining 20% for validation. The models were trained for 200 epochs. Three runs were performed with the GEX to TCR ratio varying between 1:1, 2:1 and 3:1.

Each run produced 15 trials and each trial had a different combination of model hyperparameters resulting from an automated hyperparameter grid search. The ‘best’ trial (lowest validation loss) was indicated at the end of each run, however when we manually inspected all the trial results, we found the ‘best’ trials showed strong variations between different donors. Thus, we selected one representative result from each run with minimal cross-donor batch effects for **Supplementary Fig. 4b**.

### Pseudotime inference combining ILC/NK and T cells

#### Pseudotime inference using TRBJ

scRNA-seq data of human fetal lymphoid cells from Suo et al. 2022^3^ was integrated with αβTCR data as described above. Only DN(early)_T, DN(P)_T, DN(Q)_T, DP(P)_T, DP(Q)_T, ILC2, ILC3, CYCLING_ILC, NK, CYCLING_NK cells with TRBJ were included for the trajectory analysis. Neighborhood graph (k=50) and UMAP was re-calculated using scVI latent factors similar to above.

For pseudotime trajectory analysis, *palantir*^32^ (v1.0.1) was used and a diffusion map was computed using the first five PCs. The root cell was chosen to be the neighborhood with the highest CD34 expression, and the two terminal states were chosen with the largest and smallest values on the UMAP1 axis for T and NK/ILC cell neighborhoods respectively (**Supplementary Fig. 10a**). Pseudotime and branch probabilities to the terminal states were then computed and projected back from neighborhoods (**Fig. 5b**) to cells (**Fig. 5c** top panel).

#### Gene expression trend in DN T cells along pseudotime

Chatterjee’s correlations^74^ were computed between gene expression and inferred pseudotime within DN T cells for all genes that were expressed in at least 50 cells. Chatterjee’s correlation was chosen instead of Pearson’s or Spearman’s correlation to look for any functional change and not restricted to a monotonic change. TFs^84^ and genes encoding cell surface proteins that had significantly high Chatterjee’s correlation with pseudotime (BH adjusted P-value < 0.05, and correlation coefficient > 0.1) were shown in **Fig. 5c** and

Supplementary Fig. 10b respectively.

### Code and data availability

*Dandelion* is implemented as an open-source package in Python 3 (https://github.com/zktuong/dandelion) with tutorials available at https://sc-dandelion.readthedocs.io/en/latest/. The tool and workflow is also available through an interactive online Google Colab notebook at https://colab.research.google.com/github/zktuong/dandelion/blob/master/container/dandelion_singularity.ipynb. Code and data used to generate figures and perform analyses in the manuscript are available at https://github.com/zktuong/dandelion-demo-files/dandelion_manuscript. Raw sequencing data for newly generated sequencing libraries have been deposited in ArrayExpress (accession number E-MTAB-12524).

## Acknowledgements

We acknowledge the Cellular Genetics IT, New Pipeline Group and DNA pipelines of Sanger Institute. K.B.M. and S.A.T. are supported by Wellcome (WT211276/Z/18/Z, 108413/A/15/D and Sanger core grant WT206194). K.B.M. acknowledges funding from the MRC (MR/S035907/1). M.H. is supported by Wellcome (grant WT107931/Z/15/Z), the Lister Institute for Preventive Medicine, NIHR, and the Newcastle Biomedical Research Centre. S.A.T. is supported by an ERC Consolidator Grant ThDEFINE (646794). C.S. is supported by a Wellcome Trust Ph.D. Fellowship for Clinicians. Z.K.T. and M.R.C. are supported by a Medical Research Council Research Project Grant (MR/S035842/1). M.R.C. is supported by the National Institute of Health Research (NIHR) Research Professorship (RP-2017-08-ST2-002), a Wellcome Investigator Award (220268/Z/20/Z), the Blood and Transplant Research Unit in Organ Donation and the NIHR Cambridge Biomedical Research Centre. This publication is part of the Human Cell Atlas (www.humancellatlas.org/publications). We would like to thank reviewers for their thoughtful comments and suggestions, which helped us to improve the quality of the manuscript.

## Author contributions

C.S., Z.K.T., M.R.C. and S.A.T. conceived the initial project. C.S. and Z.K.T. set up and directed the study. C.S., K.P., E.D., L.M.D. and Z.K.T. performed bioinformatic analyses. C.S., K.P. and Z.K.T developed the software. C.S. and R.V.B. performed cell culture experiments. E.D., R.G.H.L., R.V.B., R.V., M.H., K.B.M., M.R.C., and S.A.T. provided intellectual input. M.R.C. and S.A.T. acquired funding. C.S., K.P. and Z.K.T. wrote the manuscript. All authors read and/or edited the manuscript.

## Supplementary Figures

**Extended Data Fig. 1.**
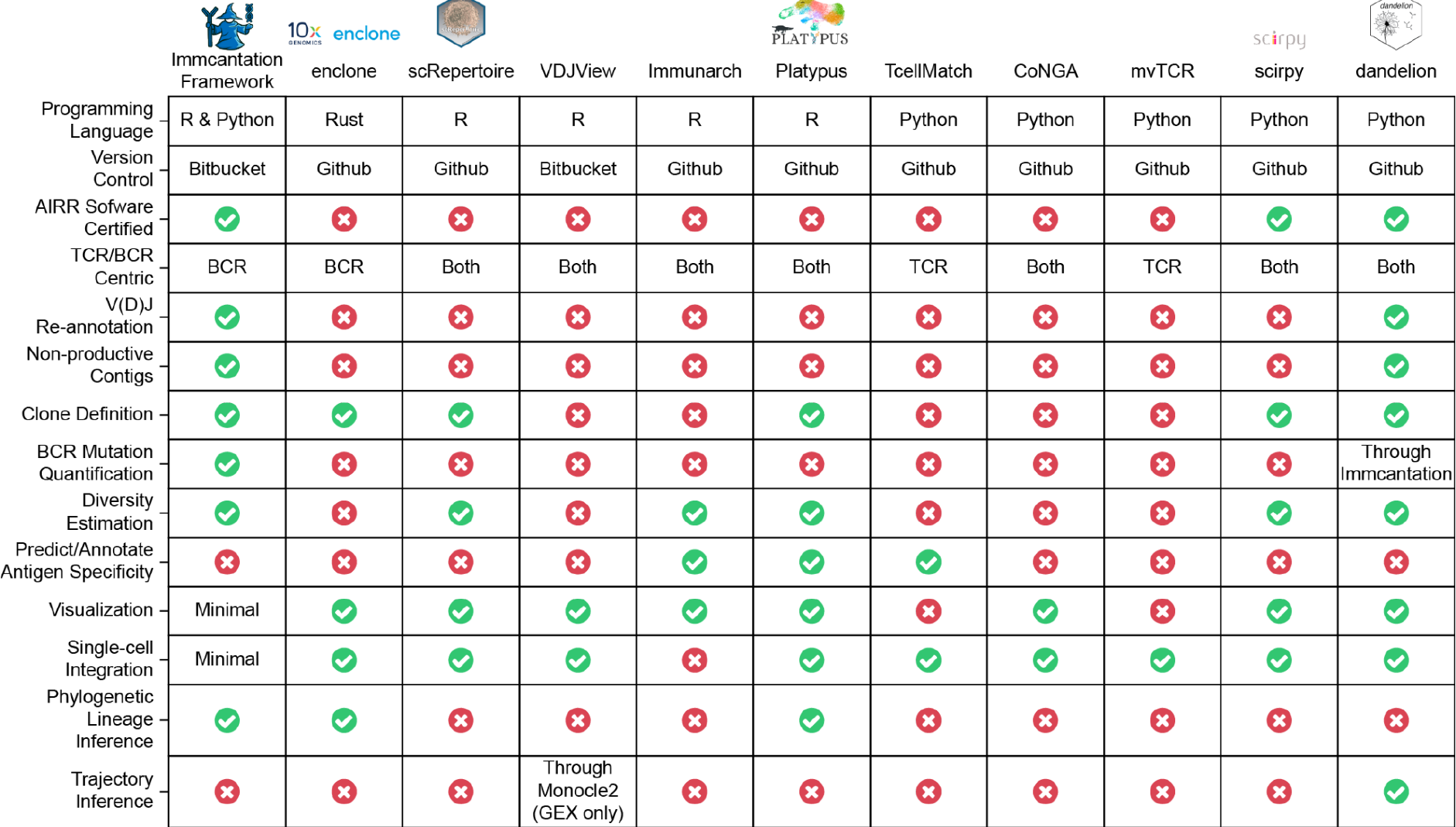
List of features included in AIR repertoire analysis pipelines. A table outlining the features of a non-exhaustive list of other methods compared to *Dandelion*. Handling of non-productive contigs (with or without V gene annotation) is not common across the various software packages. While the *Immcantation* workflow is capable of handling the data, contigs without V genes are typically diverted to a “failed” file but can be retrieved separately. The output from *Dandelion* is compatible with any AIRR-compliant softwares e.g. *Dandelion* output can be passed to *Immcantation* to perform phylogenetic lineage inference.

**Extended Data Fig. 2.**
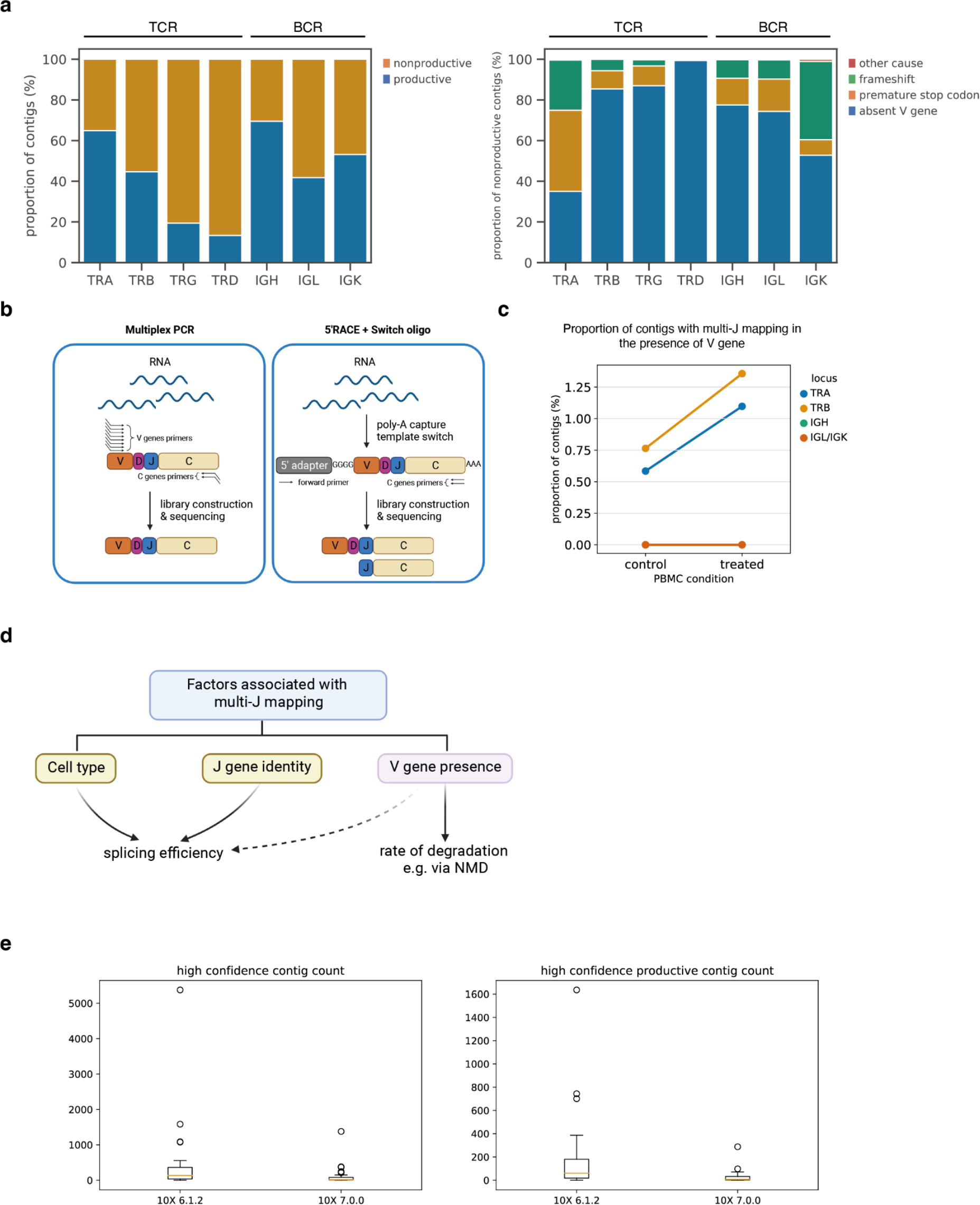
*Dandelion* offers improved contig annotations. a, Left: barplot of proportion of contigs that are productive or non-productive in each locus. Right: barplot showing the causes of non-productive contigs in each locus. For both plots, sc-γδTCR, - αβTCR and -BCR data were taken from Suo et al. 2022^3^ excluding thymus samples. **b,** Schematic illustration showing that mRNA without V genes would be captured by 5′RACE + Switch oligo technique but not by multiplex PCR strategy. **c,** Pointplot of proportion of contigs with multi-J mapping in the presence of V gene in control and cycloheximide-treated PBMC samples. Points are colored by locus of TCR/BCR. For both IGH and IGL/IGK, the proportions were 0% in control and treated. **d,** Schematic illustration showing the factors associated with multi-J mapping and the proposed mechanisms. **e,** Boxplots of sc-γδTCR contig counts annotated by 10X *cellranger vdj* v6.1.2 *versus* v7.0.0 using data from Suo et al. 2022^3^. Left: all high confidence contigs (*P*-value 5.43e-6, r 0.91 in the Wilcoxon signed-rank test). Right: high confidence productive contigs (*P*-value 1.69e-6, r 0.96 in the Wilcoxon signed-rank test).

**Extended Data Fig. 3.**
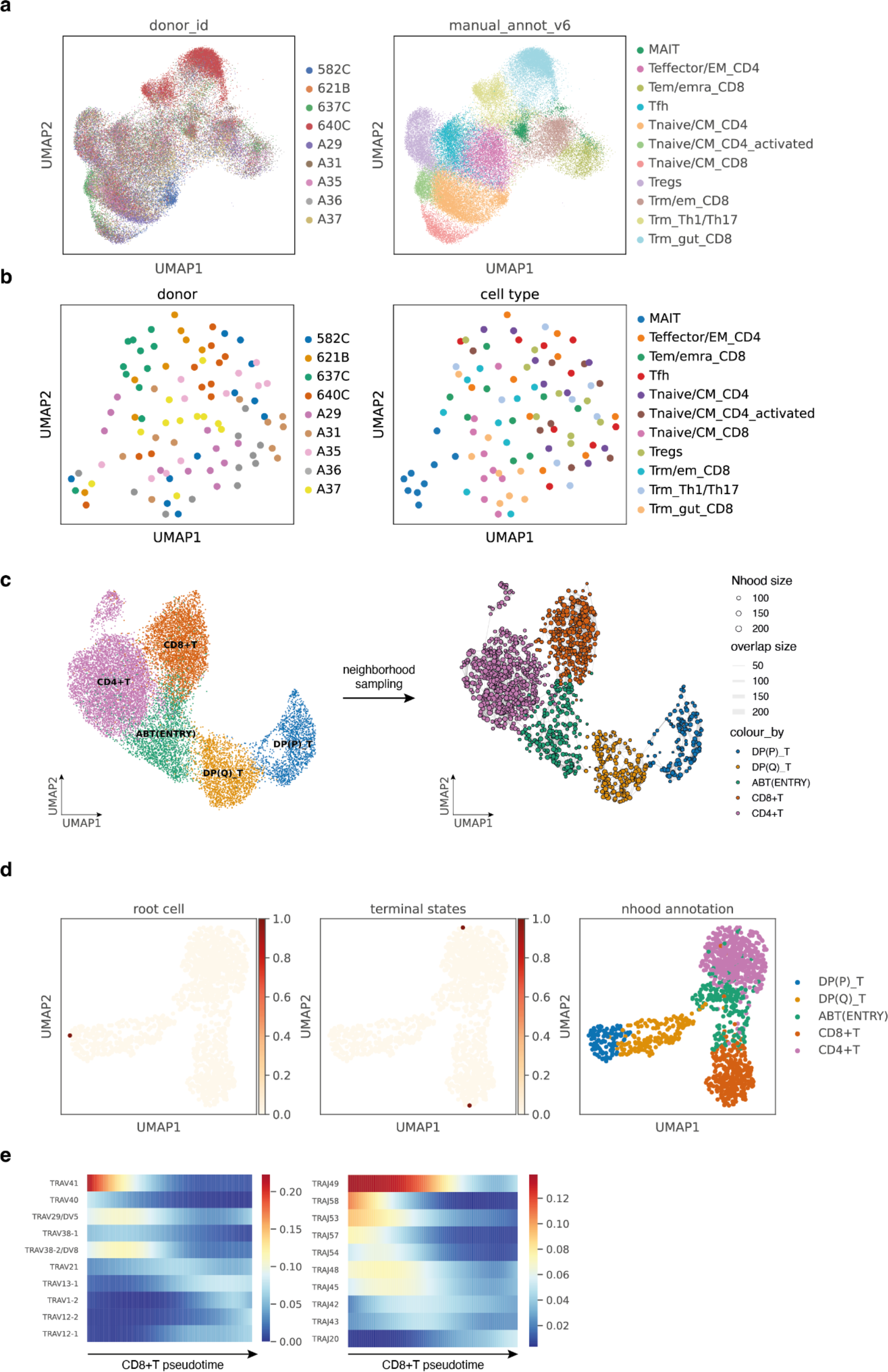
V(D)J feature space. a, Gene expression UMAP of all T cells from Conde et al. 2022^5^, colored by donor ID (left) or high-level cell type annotations (right). Each point represents a cell. **b,** UMAP of the pseudo-bulk V(D)J feature space of the same cells as in **a**, colored by donor ID (left) or high-level cell type annotations (right). Each point represents a cell pseudo-bulk. **c,** Left: UMAP of DP to mature T cells with paired productive αβTCR in data from Suo et al. 2022^3^. Each point represents a cell, colored by cell types. Right: cell neighborhood graph on the same UMAP embedding. Each point represents a cell neighborhood, colored by cell types. The point size represents neighborhood size, with connecting edges representing overlapping cell numbers between any two neighborhoods. Only edges with more than 30 overlapping cells are shown. The layout of nodes is determined by the position of the neighborhood index cell in the UMAP on the left. **d,** The root cell and terminal states selected for pseudotime inference in **Fig. 3c. e,** Gene expression trends over CD8+T pseudotime imputed with *palantir*^32^. Only the top 10 most frequently used TRAV or TRAJ genes are shown.

**Extended Data Fig. 4.**
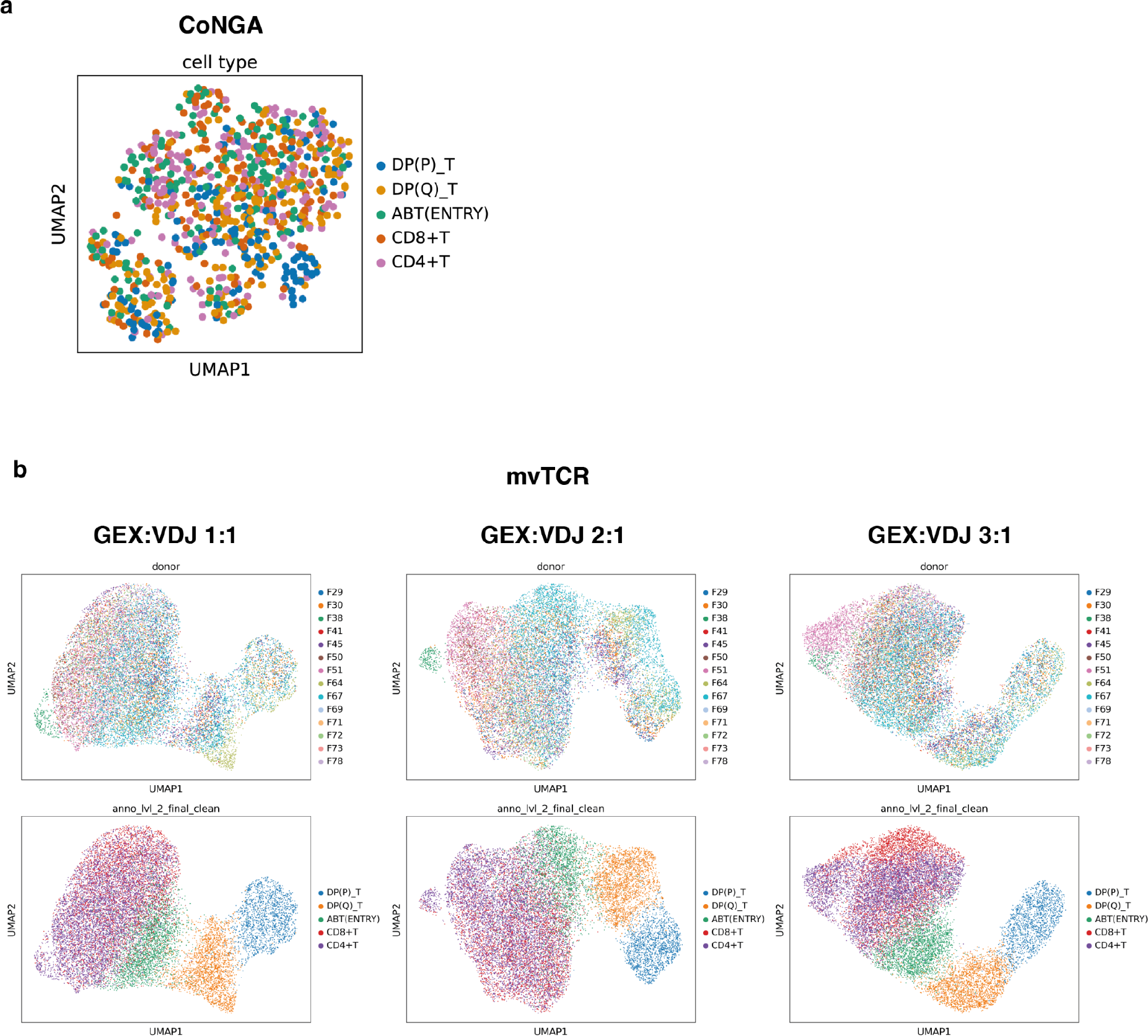
Embedding with alternative methods. a, UMAP representation of tcrdist-derived PCA coordinates of VDJ data computed by CoNGA^21^, with the same dataset as used in **Supplementary Fig. 3c**, colored by cell types. **b,** UMAP representation of joint gene expression and TCR embedding computed by mvTCR^22^ with varying weights for GEX and VDJ input, on the same dataset as used in **Supplementary Fig. 3c**. Cells are colored by donor ID (top panel) or cell types (bottom panel).

**Extended Data Fig. 5.**
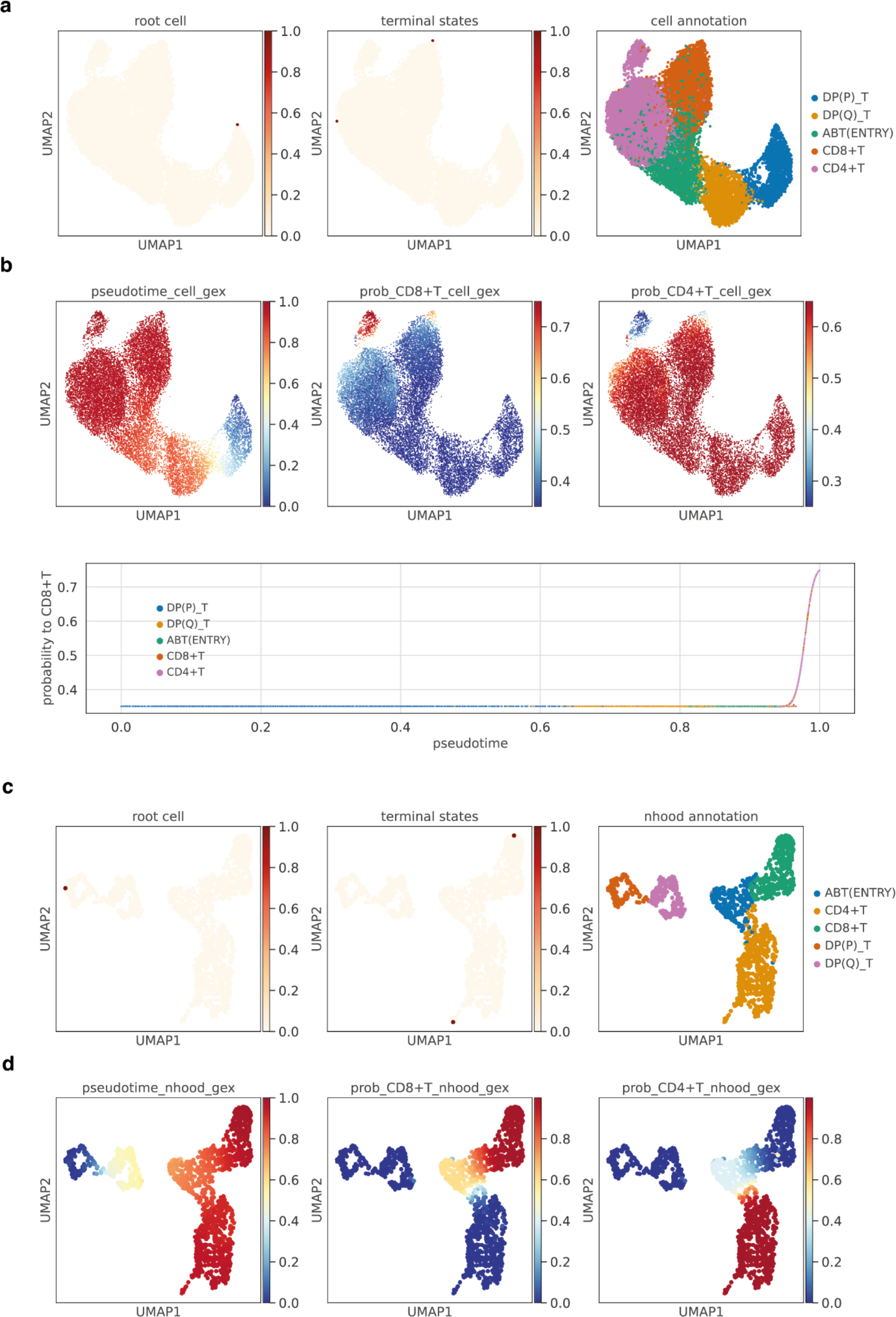
T cell development pseudotime inference comparison. a, DP to mature T cells with paired productive αβTCR in data from Suo et al. 2022^3^, on the same UMAP embedding as in **Fig. 4a** and **Supplementary Fig. 3c**. The first two panels show the root cell and terminal states selected for pseudotime inferred directly from single-cell gene expression. The last panel shows the cell types. **b,** Top: pseudotime and branch probabilities inferred directly from single-cell gene expression on the same UMAP embedding as in **a**. Bottom: scatterplot of branch probability to CD8+T against pseudotime. Each point represents a cell. **c,** UMAP of neighborhood GEX space, with the same neighborhoods as sampled in **Supplementary Fig. 3c** and UMAP embedding computed on gene expression pseudo-bulked by neighborhoods. Each point represents a cell neighborhood. The first two panels show the root cell and terminal states selected for pseudotime inferred from neighborhood GEX space. The last panel shows the cell types. **d,** Inferred pseudotime, and branch probabilities to CD8+T and to CD4+T respectively overlaid onto the same UMAP embedding in **c**.

**Extended Data Fig. 6.**
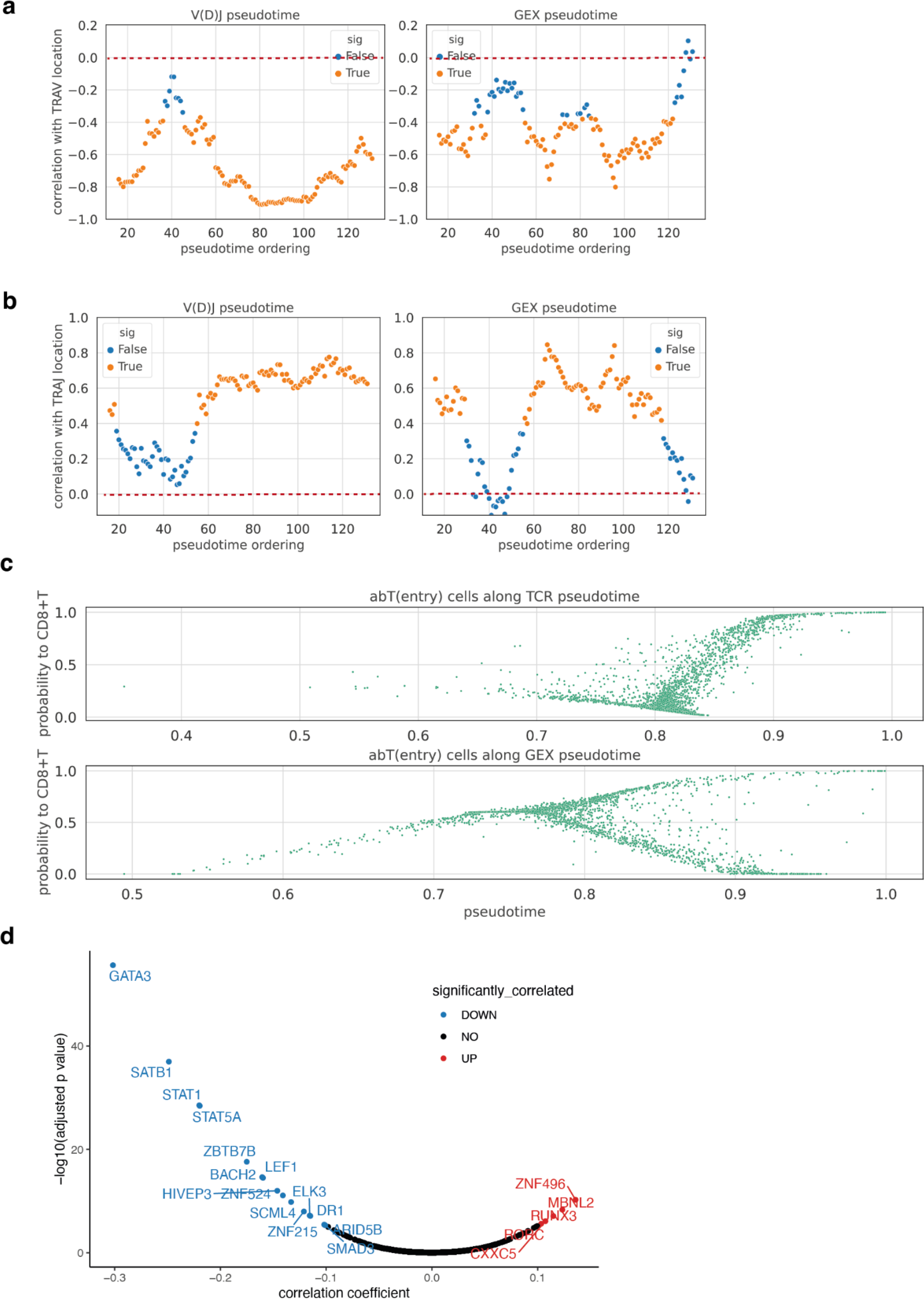
Comparing pseudotime inferred from neighborhood V(D)J space or GEX space. a, Pearson’s correlation coefficients of pseudotime order and average relative TRAV location over sliding windows of 30 adjacent neighborhoods on the pseudotime order (left: pseudotime inferred from neighborhood V(D)J space; right: pseudotime inferred from neighborhood GEX space). *Y*-axis is the correlation coefficient and the x-axis is the median pseudotime order of the 30 adjacent neighborhoods. The color of the points represents statistical significance (orange: *P*-value from the Pearson’s correlation < 0.05; blue: *P*-value ≥ 0.05). The red dashed lines mark the correlation coefficient of 0. **b,** The same plot as in **a** but for TRAJ. **c,** Scatterplots of branch probability to CD8+T against pseudotime in abT(entry) cells. Each point represents a cell. Top panel: pseudotime inferred from neighborhood V(D)J space as in **Fig. 4a** top panel. Bottom panel: pseudotime inferred from neighborhood GEX space as in **Fig. 4a** bottom right panel. **d,** Volcano plot summarizing results of TFs that are correlated with branch probabilities to CD8+T lineage in V(D)J pseudotime within abT(entry) cells. The *y*-axis is the -log10(BH adjusted *P*-value) and the *x*- axis is the correlation coefficient. Labeled TFs that had significant (BH adjusted *P*-value < 0.05) positive correlations (correlation coefficient > 0.1) were colored in red, the ones with significant negative correlations (correlation coefficient < -0.1) were colored in blue, and the rest were colored in black.

**Extended Data Fig. 7.**
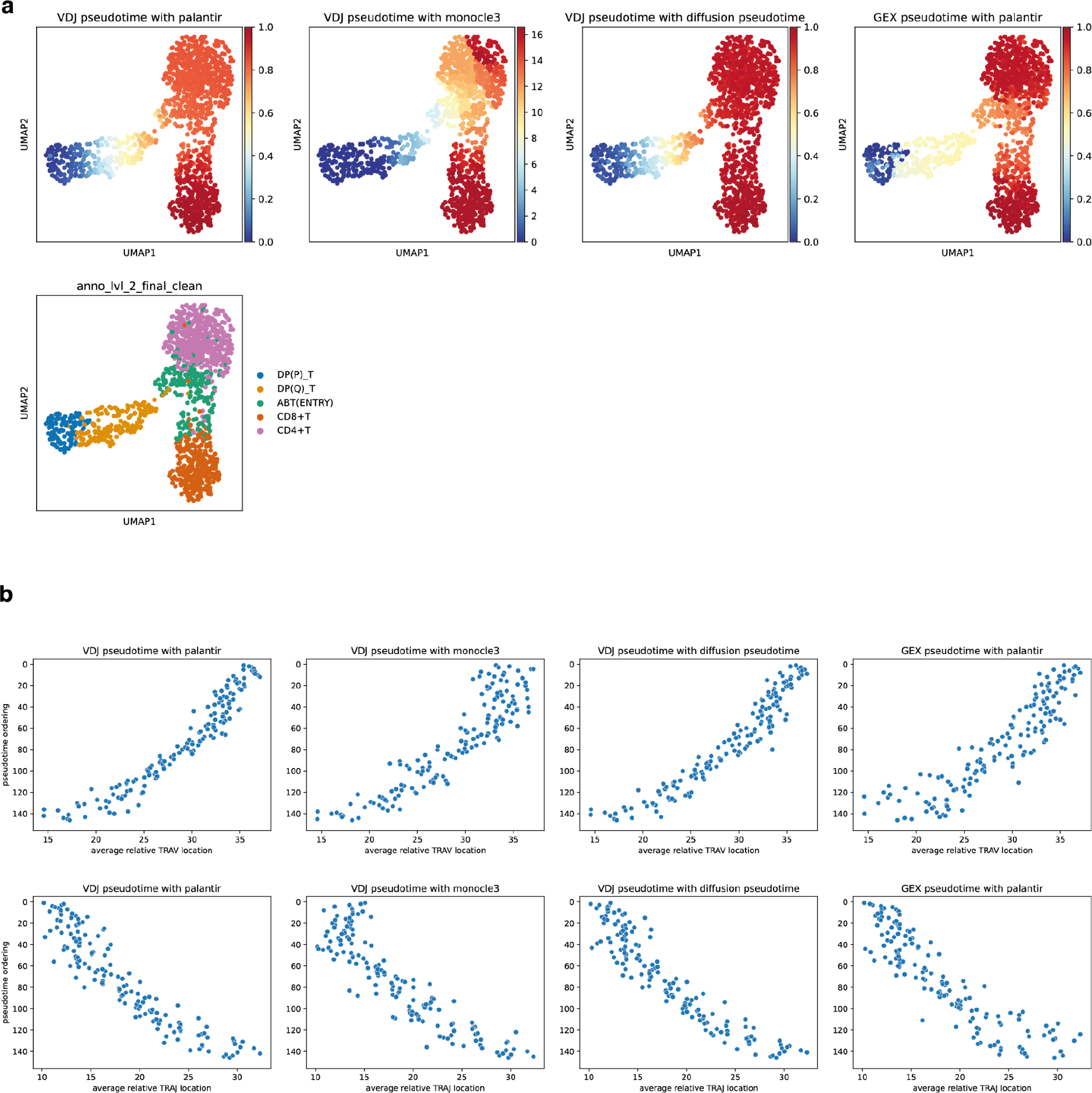
Pseudotime inferred with different trajectory inference methods. a, First three panels display pseudotime inferred from neighborhood V(D)J space using *palantir*^32^, *monocle3*^44^, and *diffusion pseudotime*^45^ respectively, overlaid onto the same UMAP embedding as in Fig. 3c with each point represents a cell neighborhood. The fourth panel represents the pseudotime inferred from neighborhood GEX space using *palantir*^32^. The last panel represents the dominant cell type in each neighborhood. **b,** Scatterplots of the pseudotime ordering against the average relative TRAV (top) and TRAJ (bottom) location. Each point represents a cell neighborhood. Each TRAV or TRAJ gene is encoded numerically for its relative genomic order. The *x*-axis represents the average TRAV/TRAJ relative location for each cell neighborhood. The *y*-axis represents the pseudotime order inferred from neighborhood V(D)J space using *palantir*^32^, *monocle3*^44^, and *diffusion pseudotime*^45^, and the pseudotime order inferred from neighborhood GEX space using *palantir*^32^ respectively. The Pearson’s correlations are -0.95, -0.91, -0.95, and -0.90 respectively (*P*-values of 4.8e-76, 4.9e-56, 2.1e-74, and 7.4e-54) for TRAV, and 0.93, 0.90, 0.93, and 0.89 respectively (*P*-values of 1.7e-62, 3.8e-54, 7.6e-65, and 4.2e-52) for TRAJ.

**Extended Data Fig. 8.**
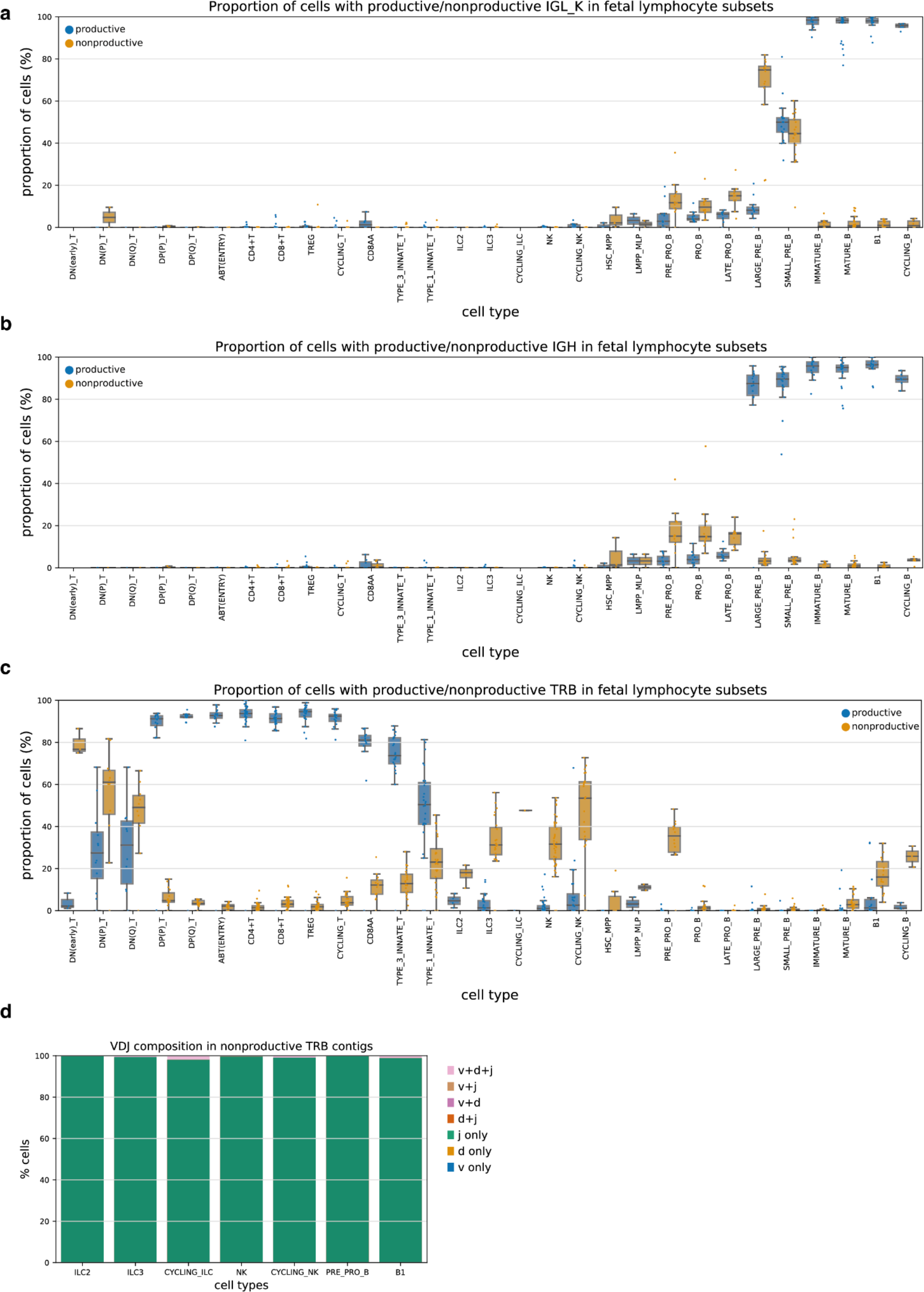
Non-productive BCR and TCR. a,b,c, Boxplot of the proportion of cells with productive (blue) or non-productive (orange) BCR light chain (**a**) and heavy chain (**b**), and TRB (**c**) in different fetal lymphocyte subsets. Each point represents a sample and data were taken from Suo et al. 2022^3^. Only samples with at least 20 cells are shown. Boxes capture the first to third quartiles and whisks span a further 1.5X interquartile range on each side of the box. **d,** Barplot showing the VDJ composition of non-productive TRB contigs in selected lymphocyte subsets from **Fig. 5a**.

**Extended Data Fig. 9.**
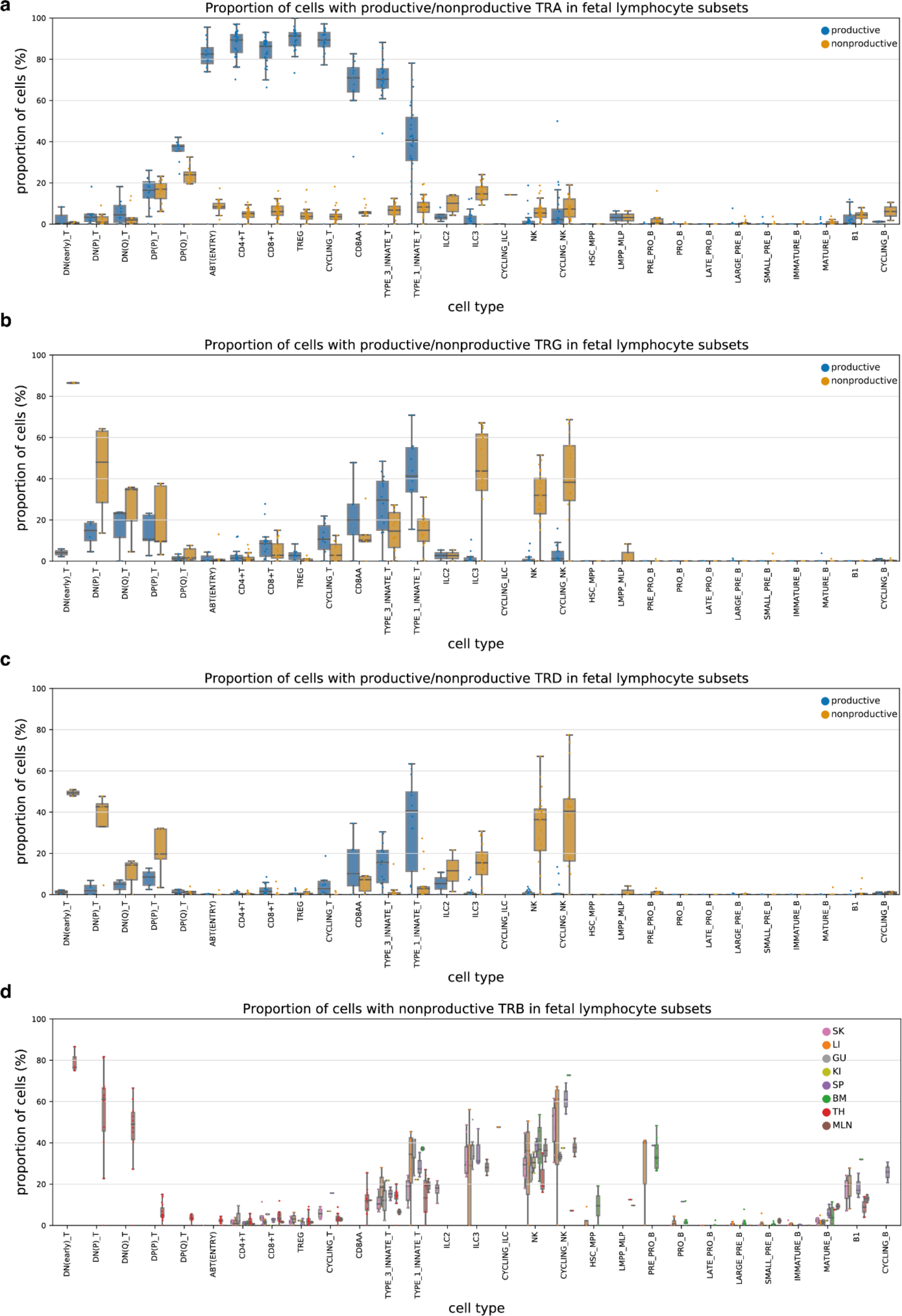
Non-productive TCR. a,b,c, Boxplot of the proportion of cells with productive (blue) or non-productive (orange) TRA (**a**), TRG (**b**) and TRD (**c**) in different fetal lymphocyte subsets. Each point represents a sample and data were taken from Suo et al. 2022^3^. Only samples with at least 20 cells are shown. Boxes capture the first to third quartiles and whisks span a further 1.5X interquartile range on each side of the box. **d,** Boxplot of the proportion of cells with non-productive TRB in different fetal lymphocyte subsets, colored by organs. Each point represents a sample. Only samples with at least 20 cells are shown. Boxes capture the first to third quartiles and whisks span a further 1.5X interquartile range on each side of the box.

**Extended Data Fig. 10.**
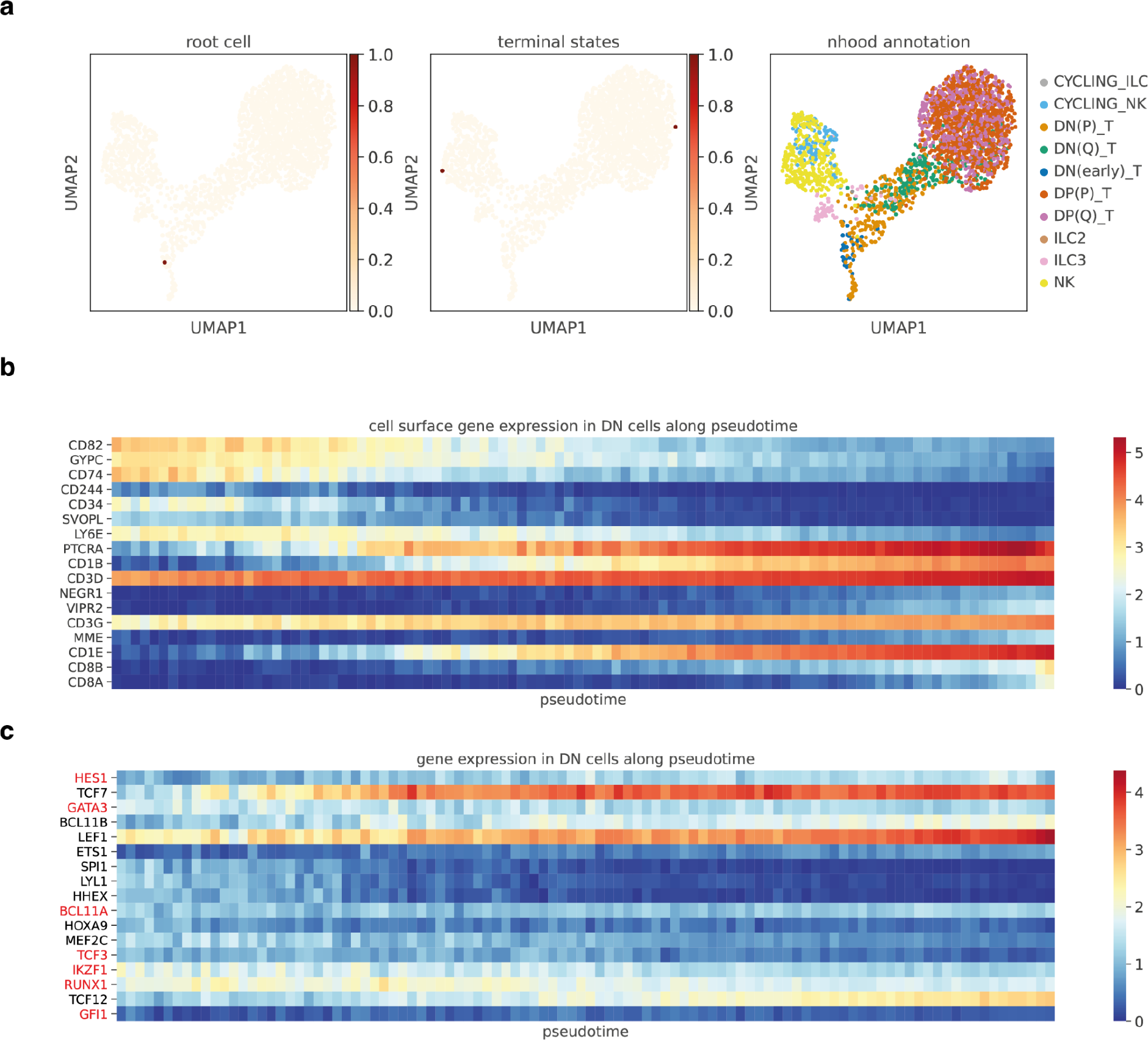
TRBJ-based trajectory for ILC/NK/T cell lineage. a, Neighborhood V(D)J feature space covering ILC, NK and developing T cells with TRBJ on the same UMAP embedding as in **Fig. 5b**. The first two panels show the root cell and terminal states selected for pseudotime inference. The last panel shows the cell types. **b,** Heatmap of gene expression for genes encoding cell surface proteins across pseudotime in DN T cells. Pseudotime is equally divided into 100 bins, and the average gene expression is calculated for DN T cells with pseudotime that falls within each bin. Genes selected here had significantly high Chatterjee’s correlation with pseudotime (BH adjusted *P*-value < 0.05, and correlation coefficient > 0.1). **c,** Heatmap of gene expression for TFs known to be important in mouse DN T cell development^53^, across pseudotime in human fetal DN T cells. TFs that showed discordant expression patterns between mouse and human are highlighted in red.

**Extended Data Fig. 11.**
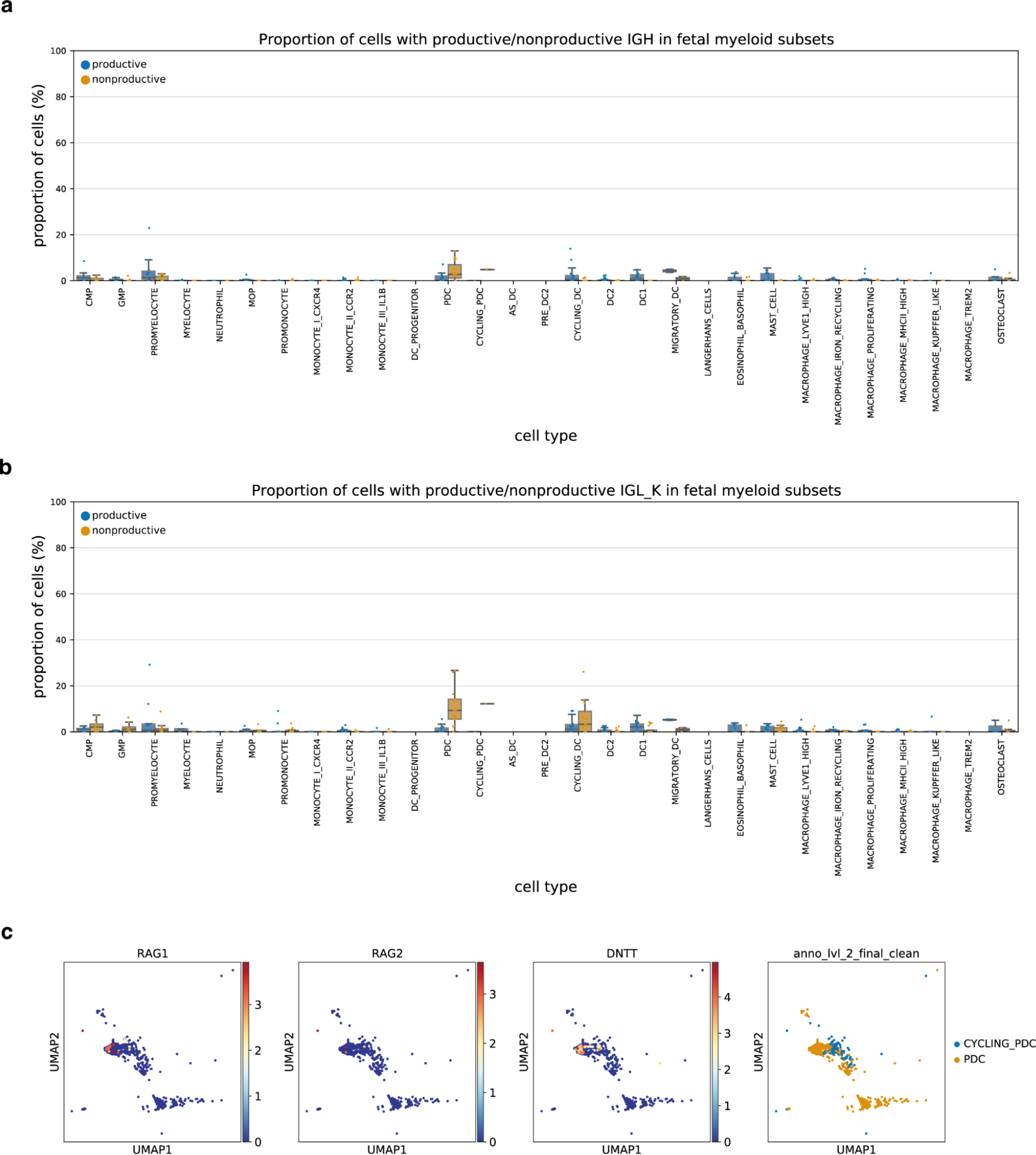
Non-productive BCR in pDC. a,b, Boxplot of the proportion of cells with productive (blue) or non-productive (orange) BCR heavy chain (**a**) and light chain (**b**) in different fetal myeloid subsets. Each point represents a sample and data were taken from Suo et al. 2022^3^. Only samples with at least 20 cells are shown. Boxes capture the first to third quartiles and whisks span a further 1.5X interquartile range on each side of the box. **c,** Expression of genes involved in V(D)J rearrangement in pDCs and cycling pDCs. Data was taken from Suo et al. 2022^3^.

## Supplementary Note 1

When *cellranger vdj* updated its contig annotation pipeline in version 3.1.0 (https://support.10xgenomics.com/single-cell-gene-expression/software/pipelines/3.1/release-notes), the priority was αβTCR and the software lost the ability to annotate γδTCR contigs.

However, the same release saw the introduction of custom enrichment primer support, which is integral for proper γδTCR contig reconstruction. γδTCR contig reannotation was reintroduced on an experimental basis in version 7.0.0 (https://support.10xgenomics.com/single-cell-gene-expression/software/pipelines/7.0/release-notes). *cellranger vdj* versions between 3.1.0 and 6.1.2 can still reconstruct γδTCR contigs, but cannot natively annotate them.

10X Genomics was aware of the issue, but it was not a priority for them as γδTCR libraries require custom enrichment primers not part of their product line-up. Early support requests would direct users to the last legacy version, 3.0.2, that supported γδTCR annotation (https://github.com/10XGenomics/cellranger/issues/45). However, this was not an ideal solution due to the lack of custom enrichment primer support. 10X Genomics subsequently revised their recommended solution to a modification of the reference, wherein all TRG sequences would be renamed to TRA and TRD would be renamed to TRB. This advice used to be available at https://kb.10xgenomics.com/hc/en-us/articles/360015793931-Can-I-detect-T-cells-with-gamma-delta-chains-in-my-V-D-J-data- but this has since been overwritten by *cellranger multi* instructions for version 7.0.0.

## Supplementary Tables

Supplementary Table 1: top_10_j_multimappers.csv (separate file)

Top 10 J gene combinations with multi-J mapping for each locus in data from Suo et al. 2022^3^, with the number of contigs containing each combination shown next to it.

Supplementary Table 2: LR_results.csv (separate file)

Logistic regression results exploring factors associated with multi-J mapping presence in data from Suo et al. 2022^3^.

Supplementary Table 3: LR_results_combined.csv (separate file)

Logistic regression results exploring factors associated with multi-J mapping presence in control and cycloheximide-treated PBMC data.

Supplementary Table 4: j_sequence_affect_j_multimapper.csv (separate file)

List of leftmost (5′ end) J genes that had significant association with increased or decreased multi-J mapping, together with the sequences of their last 10 nucleotides at 3′ ends and the first 11 nucleotides of its 3′ end intron.

Supplementary Table 5: panimmune_differential_VDJ.csv (separate file)

Differential V(D)J usage across CD4+T, CD8+T, and MAIT cells in data from Conde et al. 2022^5^.

Supplementary Table 6: abtentry_cor_result.csv (separate file)

Pearson’s correlation coefficients and BH adjusted *P*-values of all genes with branch probabilities to CD8+T lineage within abT(entry) cells.

[cor_tcr] Pearson’s correlation coefficients for pseudotime inferred from neighborhood V(D)J space

[pval_tcr] Pearson’s correlation *P*-values for pseudotime inferred from neighborhood V(D)J space

[adjp_tcr] *P*-values from pval_tcr adjusted by BH procedure

[cor_gex] Pearson’s correlation coefficients for pseudotime inferred from neighborhood GEX space

[pval_gex] Pearson’s correlation *P*-values for pseudotime inferred from neighborhood GEX space

[adjp_gex] *P*-values from pval_gex adjusted by BH procedure

## References

1. Papalexi, E. & Satija, R. Single-cell RNA sequencing to explore immune cell heterogeneity. Nat. Rev. Immunol. 18, 35–45 (2018).

2. Efremova, M., Vento-Tormo, R., Park, J.-E., Teichmann, S. A. & James, K. R. Immunology in the Era of Single-Cell Technologies. Annu. Rev. Immunol. 38, 727–757 (2020).

3. Suo, C. et al. Mapping the developing human immune system across organs. Science 376, eabo0510 (2022).

4. Stephenson, E. et al. Single-cell multi-omics analysis of the immune response in COVID-19. Nat. Med. 27, 904–916 (2021).

5. Domínguez Conde, C., et al. Cross-tissue immune cell analysis reveals tissue-specific features in humans. Science 376, eabl5197 (2022).

6. Park, J.-E. et al. A cell atlas of human thymic development defines T cell repertoire formation. Science 367, (2020).

7. Lance, C. et al. Multimodal single cell data integration challenge: results and lessons learned. bioRxiv 2022.04.11.487796 (2022) doi:10.1101/2022.04.11.487796.

8. Lee, J., Hyeon, D. Y. & Hwang, D. Single-cell multiomics: technologies and data analysis methods. Exp. Mol. Med. 52, 1428–1442 (2020).

9. Roth, D. B. V(D)J Recombination: Mechanism, Errors, and Fidelity. Microbiol Spectr 2, (2014).

10. Vander Heiden, J. A., et al. AIRR Community Standardized Representations for Annotated Immune Repertoires. Front. Immunol. 9, (2018).

11. Rubelt, F. et al. Adaptive Immune Receptor Repertoire Community recommendations for sharing immune-repertoire sequencing data. Nat. Immunol. 18, 1274–1278 (2017).

12. Breden, F. et al. Reproducibility and Reuse of Adaptive Immune Receptor Repertoire Data. Front. Immunol. 8, 1418 (2017).

13. Sturm, G. et al. Scirpy: a Scanpy extension for analyzing single-cell T-cell receptor-sequencing data. Bioinformatics 36, 4817–4818 (2020).

14. Wolf, F. A., Angerer, P. & Theis, F. J. SCANPY: large-scale single-cell gene expression data analysis. Genome Biol. 19, 15 (2018).

15. Borcherding, N., Bormann, N. L. & Kraus, G. scRepertoire: An R-based toolkit for single-cell immune receptor analysis. F1000Res. 9, 47 (2020).

16. Butler, A., Hoffman, P., Smibert, P., Papalexi, E. & Satija, R. Integrating single-cell transcriptomic data across different conditions, technologies, and species. Nat. Biotechnol. 36, 411–420 (2018).

17. Fischer, D. S., Wu, Y., Schubert, B. & Theis, F. J. Predicting antigen specificity of single T cells based on TCR CDR3 regions. Mol. Syst. Biol. 16, e9416 (2020).

18. Yermanos, A., et al. Platypus: an open-access software for integrating lymphocyte single-cell immune repertoires with transcriptomes. NAR Genom Bioinform 3, lqab023 (2021).

19. Popov, A. immunomind/immunarch: Immunarch 0.7.0. (2022). doi:10.5281/zenodo.6984421.

20. Pogorelyy, M. V. et al. Detecting T cell receptors involved in immune responses from single repertoire snapshots. PLoS Biol. 17, e3000314 (2019).

21. Schattgen, S. A. et al. Integrating T cell receptor sequences and transcriptional profiles by clonotype neighbor graph analysis (CoNGA). Nat. Biotechnol. 40, 54–63 (2022).

22. Drost, F. et al. Integrating T-cell receptor and transcriptome for large-scale single-cell immune profiling analysis. bioRxiv (2021) doi:10.1101/2021.06.24.449733.

23. Gupta, N. T. et al. Change-O: a toolkit for analyzing large-scale B cell immunoglobulin repertoire sequencing data. Bioinformatics 31, 3356–3358 (2015).

24. Virshup, I., Rybakov, S., Theis, F. J., Angerer, P. & Alexander Wolf, F. anndata: Annotated data. bioRxiv 2021.12.16.473007 (2021) doi:10.1101/2021.12.16.473007.

25. Ye, J., Ma, N., Madden, T. L. & Ostell, J. M. IgBLAST: an immunoglobulin variable domain sequence analysis tool. Nucleic Acids Res. 41, W34–40 (2013).

26. Lefranc, M. P. et al. IMGT, the international ImMunoGeneTics database. Nucleic Acids Res. 27, 209–212 (1999).

27. Le Hir, H., Gatfield, D., Izaurralde, E. & Moore, M. J. The exon–exon junction complex provides a binding platform for factors involved in mRNA export and nonsense-mediated mRNA decay. EMBO J. 20, 4987–4997 (2001).

28. Irimia, M. et al. Complex selection on 5’ splice sites in intron-rich organisms. Genome Res. 19, 2021–2027 (2009).

29. Song, L. et al. TRUST4: immune repertoire reconstruction from bulk and single-cell RNA-seq data. Nat. Methods 18, 627–630 (2021).

30. Dann, E., Henderson, N. C., Teichmann, S. A., Morgan, M. D. & Marioni, J. C. Differential abundance testing on single-cell data using k-nearest neighbor graphs. Nat. Biotechnol. 40, 245– 253 (2022).

31. Saelens, W., Cannoodt, R., Todorov, H. & Saeys, Y. A comparison of single-cell trajectory inference methods. Nat. Biotechnol. 37, 547–554 (2019).

32. Setty, M. et al. Characterization of cell fate probabilities in single-cell data with Palantir. Nat. Biotechnol. 37, 451–460 (2019).

33. Carico, Z. M., Roy Choudhury, K., Zhang, B., Zhuang, Y. & Krangel, M. S. Tcrd Rearrangement Redirects a Processive Tcra Recombination Program to Expand the Tcra Repertoire. Cell Rep. 19, 2157–2173 (2017).

34. Singer, A., Adoro, S. & Park, J.-H. Lineage fate and intense debate: myths, models and mechanisms of CD4- versus CD8-lineage choice. Nat. Rev. Immunol. 8, 788–801 (2008).

35. Karimi, M. M. et al. The order and logic of CD4 versus CD8 lineage choice and differentiation in mouse thymus. Nat. Commun. 12, 1–14 (2021).

36. Kirchner, J. & Bevan, M. J. ITM2A is induced during thymocyte selection and T cell activation and causes downregulation of CD8 when overexpressed in CD4(+)CD8(+) double positive thymocytes. J. Exp. Med. 190, 217–228 (1999).

37. Taniuchi, I. et al. Differential requirements for Runx proteins in CD4 repression and epigenetic silencing during T lymphocyte development. Cell 111, 621–633 (2002).

38. Sato, T. et al. Dual functions of Runx proteins for reactivating CD8 and silencing CD4 at the commitment process into CD8 thymocytes. Immunity 22, 317–328 (2005).

39. He, X. et al. The zinc finger transcription factor Th-POK regulates CD4 versus CD8 T-cell lineage commitment. Nature 433, 826–833 (2005).

40. Sun, G. et al. The zinc finger protein cKrox directs CD4 lineage differentiation during intrathymic T cell positive selection. Nat. Immunol. 6, 373–381 (2005).

41. Aliahmad, P. & Kaye, J. Development of all CD4 T lineages requires nuclear factor TOX. J. Exp. Med. 205, 245–256 (2008).

42. Hernández-Hoyos, G., Anderson, M. K., Wang, C., Rothenberg, E. V. & Alberola-Ila, J. GATA- 3 expression is controlled by TCR signals and regulates CD4/CD8 differentiation. Immunity 19, 83–94 (2003).

43. Pai, S.-Y. et al. Critical roles for transcription factor GATA-3 in thymocyte development. Immunity 19, 863–875 (2003).

44. Cao, J. et al. The single-cell transcriptional landscape of mammalian organogenesis. Nature 566, 496–502 (2019).

45. Haghverdi, L., Büttner, M., Wolf, F. A., Buettner, F. & Theis, F. J. Diffusion pseudotime robustly reconstructs lineage branching. Nat. Methods 13, 845–848 (2016).

46. Clark, M. R., Mandal, M., Ochiai, K. & Singh, H. Orchestrating B cell lymphopoiesis through interplay of IL-7 receptor and pre-B cell receptor signalling. Nat. Rev. Immunol. 14, 69–80 (2014).

47. Wong, J. B. et al. B-1a cells acquire their unique characteristics by bypassing the pre-BCR selection stage. Nat. Commun. 10, 4768 (2019).

48. Kitamura, D. et al. A critical role of λ5 protein in B cell development. Cell 69, 823–831 (1992).

49. O’Byrne, S. et al. Discovery of a CD10-negative B-progenitor in human fetal life identifies unique ontogeny-related developmental programs. Blood 134, 1059–1071 (2019).

50. Shin, S. B. et al. Abortive γδTCR rearrangements suggest ILC2s are derived from T-cell precursors. Blood Adv 4, 5362–5372 (2020).

51. Qian, L., et al. Suppression of ILC2 differentiation from committed T cell precursors by E protein transcription factors. Journal of Experimental Medicine vol. 216 884–899 Preprint at https://doi.org/10.1084/jem.20182100 (2019).

52. Shin, S. B. & McNagny, K. M. ILC-You in the Thymus: A Fresh Look at Innate Lymphoid Cell Development. Front. Immunol. 12, 681110 (2021).

53. Hosokawa, H. & Rothenberg, E. V. How transcription factors drive choice of the T cell fate. Nat. Rev. Immunol. 21, 162–176 (2021).

54. Musumeci, A., Lutz, K., Winheim, E. & Krug, A. B. What Makes a pDC: Recent Advances in Understanding Plasmacytoid DC Development and Heterogeneity. Front. Immunol. 10, 1222 (2019).

55. Popescu, D.-M. et al. Decoding human fetal liver haematopoiesis. Nature 574, 365–371 (2019).

56. Corcoran, L. et al. The lymphoid past of mouse plasmacytoid cells and thymic dendritic cells. J. Immunol. 170, 4926–4932 (2003).

57. Shigematsu, H. et al. Plasmacytoid dendritic cells activate lymphoid-specific genetic programs irrespective of their cellular origin. Immunity 21, 43–53 (2004).

58. Pelayo, R. et al. Derivation of 2 categories of plasmacytoid dendritic cells in murine bone marrow. Blood 105, 4407–4415 (2005).

59. Sathe, P., Vremec, D., Wu, L., Corcoran, L. & Shortman, K. Convergent differentiation: myeloid and lymphoid pathways to murine plasmacytoid dendritic cells. Blood 121, 11–19 (2013).

60. Mak, T. W. & Saunders, M. E. The immune response. Part I: Basic Immunology 373–401 (2006).

61. Charles, A., Janeway, J., Travers, P. & Walport, M. Immunobiology: the immune system in health and disease. Current Biology Ltd./Garland.

62. Elhanati, Y. et al. Inferring processes underlying B-cell repertoire diversity. Philos. Trans. R. Soc. Lond. B Biol. Sci. 370, (2015).

63. Sethna, Z. et al. Population variability in the generation and selection of T-cell repertoires. PLoS Comput. Biol. 16, e1008394 (2020).

64. Okoreeh, M. K., et al. Asymmetrical forward and reverse developmental trajectories determine molecular programs of B cell antigen receptor editing. Sci Immunol 7, eabm1664 (2022).

65. Jaffe, D. B. et al. Functional antibodies exhibit light chain coherence. Nature (2022) doi:10.1038/s41586-022-05371-z.

66. Montecino-Rodriguez, E. & Dorshkind, K. B-1 B cell development in the fetus and adult. Immunity 36, 13–21 (2012).

67. Herzenberg, L. A. & Herzenberg, L. A. Toward a layered immune system. Cell 59, 953–954 (1989).

68. Solvason, N., Lehuen, A. & Kearney, J. F. An embryonic source of Ly1 but not conventional B cells. Int. Immunol. 3, 543–550 (1991).

69. Montecino-Rodriguez, E., Leathers, H. & Dorshkind, K. Identification of a B-1 B cell-specified progenitor. Nat. Immunol. 7, 293–301 (2006).

70. Esplin, B. L., Welner, R. S., Zhang, Q., Borghesi, L. A. & Kincade, P. W. A differentiation pathway for B1 cells in adult bone marrow. Proc. Natl. Acad. Sci. U. S. A. 106, 5773–5778 (2009).

71. Yoshimoto, M. et al. Embryonic day 9 yolk sac and intra-embryonic hemogenic endothelium independently generate a B-1 and marginal zone progenitor lacking B-2 potential. Proc. Natl. Acad. Sci. U. S. A. 108, 1468–1473 (2011).

72. Kreslavsky, T., Wong, J. B., Fischer, M., Skok, J. A. & Busslinger, M. Control of B-1a cell development by instructive BCR signaling. Curr. Opin. Immunol. 51, 24–31 (2018).

73. Graf, R. et al. BCR-dependent lineage plasticity in mature B cells. Science 363, 748–753 (2019).

74. Chatterjee, S. A New Coefficient of Correlation. J. Am. Stat. Assoc. 116, 2009–2022 (2021).

75. Gadala-Maria, D., Yaari, G., Uduman, M. & Kleinstein, S. H. Automated analysis of high- throughput B-cell sequencing data reveals a high frequency of novel immunoglobulin V gene segment alleles. Proc. Natl. Acad. Sci. U. S. A. 112, E862–70 (2015).

76. Sleckman, B. P., Khor, B., Monroe, R. & Alt, F. W. Assembly of productive T cell receptor delta variable region genes exhibits allelic inclusion. J. Exp. Med. 188, 1465–1471 (1998).

77. Hu, Y. Efficient, high-quality force-directed graph drawing. Mathematica journal (2005).

78. Peixoto, T. P. The graph-tool python library. (2017) doi:10.6084/M9.FIGSHARE.1164194.

79. Wolock, S. L., Lopez, R. & Klein, A. M. Scrublet: Computational Identification of Cell Doublets in Single-Cell Transcriptomic Data. Cell Syst 8, 281–291.e9 (2019).

80. Benjamini, Y. & Hochberg, Y. Controlling the false discovery rate: A practical and powerful approach to multiple testing. J. R. Stat. Soc. 57, 289–300 (1995).

81. Crooks, G. E., Hon, G., Chandonia, J.-M. & Brenner, S. E. WebLogo: a sequence logo generator. Genome Res. 14, 1188–1190 (2004).

82. Kerby, D. S. The Simple Difference Formula: An Approach to Teaching Nonparametric Correlation. Comprehensive Psychology 3, 11.IT.3.1 (2014).

83. Lopez, R., Regier, J., Cole, M. B., Jordan, M. I. & Yosef, N. Deep generative modeling for single-cell transcriptomics. Nat. Methods 15, 1053–1058 (2018).

84. Lambert, S. A. et al. The Human Transcription Factors. Cell 175, 598–599 (2018).

